# Leveraging microfluidic dielectrophoresis to distinguish compositional variations of lipopolysaccharide in *E. coli*

**DOI:** 10.1101/2022.08.19.504570

**Authors:** Qianru Wang, Hyungseok Kim, Tiffany M. Halvorsen, Sijie Chen, Christopher S. Hayes, Cullen R. Buie

## Abstract

Lipopolysaccharide (LPS) is the unique feature that composes the outer leaflet of the Gram-negative bacterial cell envelope. Variations in LPS structures affect a number of physiological processes, including outer membrane permeability, antimicrobial resistance, recognition by the host immune system, biofilm formation, and interbacterial competition. Rapid characterization of LPS properties is crucial for studying the relationship between these LPS structural changes and bacterial physiology. However, current assessments of LPS structures require LPS extraction and purification followed by cumbersome proteomic analysis. This paper demonstrates one of the first high-throughput and noninvasive strategies to directly distinguish *Escherichia coli* with different LPS structures. Using a combination of three-dimensional insulator-based dielectrophoresis (3DiDEP) and cell tracking in a linear electrokinetics assay, we elucidate the effect of structural changes in *E. coli* LPS oligosaccharides on electrokinetic mobility and polarizability. We show that our platform is sufficiently sensitive to detect LPS structural variations at the molecular level. To correlate electrokinetic properties of LPS with the outer membrane permeability, we further examined effects of LPS structural variations on bacterial susceptibility to colistin, an antibiotic known to disrupt the outer membrane by targeting LPS. Our results suggest that microfluidic electrokinetic platforms employing 3DiDEP can be a useful tool for isolating and selecting bacteria based on their LPS glycoforms. Future iterations of these platforms could be leveraged for rapid profiling of pathogens based on their surface LPS structural identity.

## 1 Introduction

Lipopolysaccharide (LPS) (Raetz and Whitfield, 2002) is a glycolipid and the distinguishing feature of the outer membrane of Gram-negative bacteria, comprising over 75% of the cell surface in *E. coli* and *Salmonella* (Klein and Raina, 2019; Simpson and Trent, 2019). LPS forms the outer leaflet of the outer membrane and is an efficient permeability barrier to compounds such as antimicrobial peptides (AMPs). The chemical and structural diversity of LPS molecules is critical for the survival of pathogenic bacteria, which live similarly diverse lifestyles (Caroff and Novikov, 2019). Indeed, many bacteria have evolved the ability to modify LPS in response to changes in environmental conditions, which leads to adaptive changes in outer membrane permeability, antibiotic resistance, and acts as a means to evade the human immune response (Needham and Trent, 2013; Caroff and Novikov, 2019; Klein and Raina, 2019; Simpson and Trent, 2019). Importantly, by studying LPS structural changes and their impact on the pathogenic lifestyle, new therapeutic treatments targeting LPS modifying enzymes have been proposed (Harris et al., 2014).

LPS consists of three regions: lipid A, the core oligosaccharide, and a distal polysaccharide exhibiting the most inter-strain heterogeneity (termed O-antigen, (Caroff and Novikov, 2019)) (Figure 1a). Lipid A is an amphipathic lipid consisting of a hexa-acylated glucosamine diphosphate headgroup that is *bis*-phosphorylated in *E. coli*. The phosphate substituents of lipid A mediate salt bridge formation with divalent cations (Ca^2+^ and Mg^2+^), which are crucial to stabilize the outer membrane (Needham and Trent, 2013; Caroff and Novikov, 2019). Attached to lipid A via a 3-deoxy-D-*manno*-oct-2-ulosonic acid (Kdo) linkage is the core oligosaccharide, which can be further divided into the inner and outer core regions. In *E. coli*, the inner core region contains Kdo and heptose (Yethon et al., 1998; Wang et al., 2015; Caroff and Novikov, 2019). Due to the presence of phosphate and carboxyl groups decorating these residues, the inner core is highly negatively charged. The outer core contains glucose and heptose (Wang et al., 2015) and displays more variations in sugar composition and arrangement. Attached to the outer core is the more structurally diverse O-antigen, a long carbohydrate polymer built from a modal distribution of repeating short oligosaccharide monomers (Lerouge and Vanderleyden, 2002; Caroff and Novikov, 2019). O-antigen plays important roles in virulence and resistance to AMPs (Kim and Slauch, 1999; Lerouge and Vanderleyden, 2002; Bogomolnaya et al., 2008).

**Figure 1.**
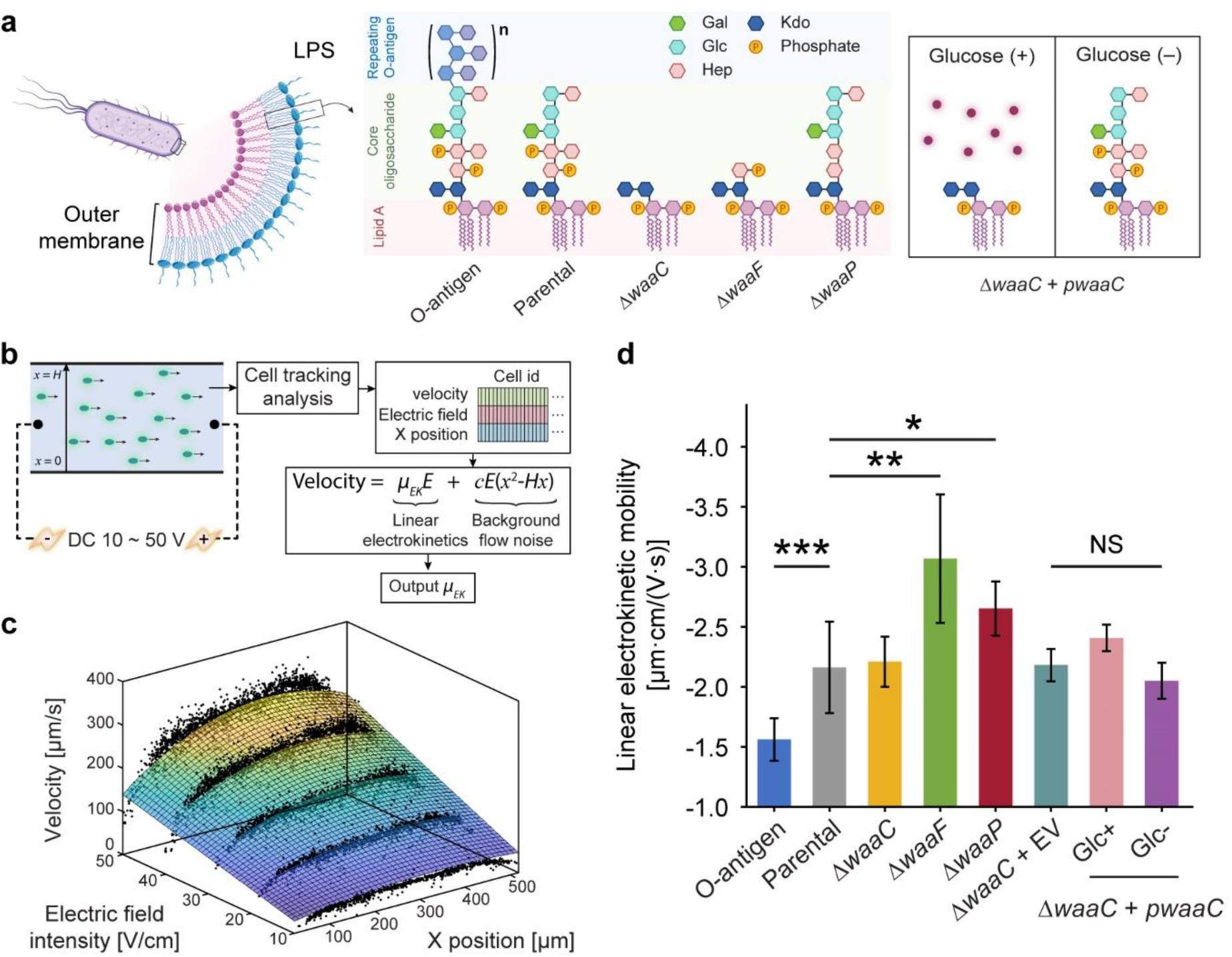
Truncating *E. coli* LPS core components affects cell electrokinetic mobility. (a) Schematic of the expected LPS structures of *E. coli* strains examined in this study. The O-antigen expressing strain is a K-12 strain with restored O16 polysaccharide synthesis. The parental strain expresses full-length LPS without O-antigen. Δ*waaC* and Δ*waaF* mutants exhibit a shorter LPS, leaving only the Kdo2-lipid A and Hep-Kdo2-lipid A moieties of the inner core, respectively. The Δ*waaP* strain lacks phosphoryl substituents on heptose (Hep) residues I and II, and is deficient in adding heptose III. The *waaC*-complementary strain (Δ*waaC* + *pwaaC*) carries the plasmid pCH450-*waaC*, whose expression is suppressed in the presence of glucose (Halvorsen et al., 2021). Abbreviations: Kdo, 3-deoxy-D-*manno*-oct-2-ulosonic acid; Hep, L-glycero-D-*manno*-heptose; Glc, glucose; Gal, galactose. (b) Workflow of the cell tracking analysis for measuring cell linear electrokinetic mobility. Cell motion in a straight PMMA channel was imaged to extract cell velocity, *x* position, and corresponding electric field, which were then used to derive linear electrokinetic mobilities (*μEK*) for individual cells. A 3D fitting model was used to exclude background flow noise (e.g. pressure-driven or electrothermal flows). (c) A representative fitting surface (color scale indicates velocity magnitudes) obtained from single cell measurements (black dots). (d) Linear electrokinetic mobilities measured for *E. coli* strains. See Supplementary Figure 7 for details comparing two approaches for the measurement. Asterisks and error bars indicate statistical difference (\**p* < 0.02, \*\**p* < 0.005, and \*\*\**p* < 0.001, NS: not significant, Kruskal-Wallis test) and standard deviations, respectively.

Genetically induced chemical modifications can occur within each region of the LPS molecule to cope with environmental stressors, and are key to the survival and persistence of pathogens in human hosts (Caroff and Novikov, 2019). A more complete understanding of these adaptive cell envelope changes would aid in therapeutic development, because LPS alterations affect small molecular uptake. For example, phosphorylation and acylation of lipid A are capable of modulating outer membrane permeability (Gunn, 2001; Raetz et al., 2007; Simpson and Trent, 2019). Moreover, addition of charged moieties or removal of sugars from the core oligosaccharide drives resistance to cationic AMPs by disrupting electrostatically-driven membrane association (Gunn, 2001; Reynolds et al., 2005; Wang et al., 2015). Finally, a large diversity of O-antigen modifications lead to inter- and intra-strain LPS compositional variations, which is not only an indicator of virulence, but also arms the pathogens by facilitating colonization and bypassing host defense mechanisms during infection (Lerouge and Vanderleyden, 2002; Bogomolnaya et al., 2008; Simpson and Trent, 2019).

It is important to understand LPS structural versatility because its composition drives AMP resistance and evasion of the host immune response (Gunn, 2001; Lerouge and Vanderleyden, 2002; Needham and Trent, 2013). However, a better understanding of LPS structural adaptability is hindered by the lack of efficient LPS phenotyping approaches. Conventional methods to characterize LPS, such as gel electrophoresis (Tsai and Frasch, 1982) and mass spectrometry (Kilár et al., 2013), require prerequisite steps for LPS extraction and purification from cell lysates, which is time-consuming, laborious, and requires use of hazardous reagents and large sample volumes. Microfluidics is a powerful tool to overcome this issue as it enables accurate and high-throughput manipulation of small sample volumes. Previous attempts to characterize LPS with microfluidics have used platforms functionalized with bio-recognition ligands, such as antibodies (Tokel et al., 2015), aptamers (Niu et al., 2019) or AMPs (Mannoor et al., 2010; Yoo et al., 2014), which bind specifically to LPS to enable pathotyping. In addition, microchip electrophoresis has been applied to LPS structural analysis by facilitating LPS detection via fluorescent dye conjugation (Kilár et al., 2008; Makszin et al., 2012). However, reproducibility of these approaches relies on high-purity and repeatable LPS extraction and separation. Although a variety of LPS extraction protocols have been developed (Wang et al., 2010), each method is limited by size bias. For instance, the phenol–water extraction method (Hickman and Ashwell, 1966) favors smooth LPS over rough LPS, whereas the ether extraction method (Galanos et al., 1969) favors the opposite. In addition, many of these LPS extraction methods suffer from impurities, low yield, and contamination with proteins and nucleic acids, which impede reliable application of the end product to sensitive downstream analysis, such as molecular and immunological experiments. Moreover, these methods often involve LPS staining steps, which are time-consuming, toxic, expensive, and likely to induce chemical modifications of LPS (e.g. silver staining) (Wang et al., 2010).

Microfluidic dielectrophoresis (DEP) (Hawkins et al., 2007; Braff et al., 2012; Jones et al., 2015; Lapizco-Encinas, 2019) systems using DC electric fields allow studies of cell surface properties exclusively (Braff et al., 2013; Jones et al., 2015; Crowther et al., 2019; Wang et al., 2019), and have demonstrated robust efficacy in separating bacteria based on a variety of surface-related properties, including surface chemical modifications (Crowther et al., 2019), biofilm formation capabilities (Braff et al., 2013), and antibiotic resistance (Jones et al., 2015). Our previous work has shown that three-dimensional insulator-based DEP (3DiDEP)(Braff et al., 2012, 2013; Wang et al., 2019) provides sensitive and effective distinction of bacteria with differential expression of *c*-type outer-membrane cytochromes, which are known to be responsible for extracellular electron transfer across the cell envelope, in electrochemically active microorganisms (Wang et al., 2019). Our technique offers sub-strain level resolution at low applied voltages, which diminishes confounding secondary flow effects, such as Joule heating and induced charge electroosmosis (Wang et al., 2017). In addition to DEP, linear electrokinetic behaviors of bacteria are also affected by cell surface appendages. Duval and Ohshima proposed an electrokinetic theory of diffuse soft particles, in which the charged cell surface components, such as LPS or outer membrane proteins, form a soft polyelectrolyte layer around the cell (Duval and Ohshima, 2006; Ohshima, 2013). Charge conditions and the spatial arrangement of polyelectrolyte segments in the soft layer substantially affect cell electrokinetics, suggesting that whole-cell electrokinetic mobility can be a biomarker to distinguish LPS structural variations. This model has been applied to distinguish *E. coli* producing different fimbriae and pilus (Francius et al., 2011) and *Acidithiobacillus ferrooxidans* grown with different metal ions (Chandraprabha et al., 2009).

In this work, we demonstrate the efficacy of 3DiDEP as a high-throughput and noninvasive approach to probe bacterial LPS structural modifications. Using 3DiDEP and linear electrokinetics, we assayed several *E. coli* K-12 strains deficient in synthesizing LPS core components, in addition to a parental strain as a wild type reference and an O-antigen expressing strain. We show that compositional variations in LPS structures drive changes in electrokinetic properties of the cell surface (i.e. polarizability and linear electrokinetic mobility), resulting in a measurable metric for predicting cell surface properties. With further investigations on colistin-mediated outer membrane permeabilization, we observe a distinct correlation between the electrokinetic properties of *E. coli* cells and their susceptibility to colistin, with colistin sensitivity increasing with cell polarizability. We demonstrate that cell surface polarizability and electrokinetic mobility can be used as a proxy for selecting bacteria based on their LPS structures in DEP-based systems, which will open a broader avenue for downstream molecular and proteomic analysis to elucidate mechanisms underlying LPS modifications and adaptive resistance to AMPs.

## 2 Materials and methods

### 2.1 LPS mutation and plasmid construction

*E. coli* K-12 LPS *waa* mutations were constructed as described previously (Halvorsen et al., 2021). Briefly, the *waaC* and *waaP* mutant strains were generated by bacteriophage λ Red-mediated recombineering: a kanamycin-resistance cassette flanked by homology arms for the up- and downstream regions of each gene was first cloned into an intermediate plasmid, then PCR amplified to generate a linear fragment for recombination. To generate the *waaF* mutant strain, a linear fragment containing a kanamycin resistance gene flanked by homology arms was PCR amplified from the Δ*waaF* Keio collection deletion strain (Baba et al., 2006) and recombineered as described above. To create an arabinose-inducible *waaC* complemented strain, the *waaC* mutant was transformed with plasmid pCH450-*waaC* (denoted by strain Δ*waaC* + *pwaaC* as a shorthand) or an empty plasmid (pCH450; strain Δ*waaC* + EV). Expression from these plasmids is suppressed in the presence of glucose. Because induction with arabinose caused a growth defect in the *waaC* complemented strain but removing glucose was sufficient to complement the *waaC* deletion, we chose to use the latter to complement.

### 2.2 Bacterial strains

Bacterial strains and plasmids used in this study are listed in Supplementary Table 1. Following the previous study constructing these strains (Halvorsen et al., 2021), all bacterial cells were inoculated from frozen stocks, and single colonies were grown in LB at 37°C overnight, then diluted 1:100 in fresh media and grown to mid-log phase before harvesting for 3DiDEP analysis. Where appropriate, media were supplemented with antibiotics at the following concentrations: kanamycin (Kan), 50 μg/mL (Sigma-Aldrich); streptomycin (Str), 50 μg/mL (Sigma-Aldrich); spectinomycin (Spm), 100 μg/mL (Sigma-Aldrich); and tetracycline (Tet), 15 μg/mL (Sigma-Aldrich). To achieve a dynamic control of *waaC* expression in the *waaC*-complementary strain (Δ*waaC* + *pwaaC*), both strain Δ*waaC +* EV (control for plasmid burden) and strain Δ*waaC* + *pwaaC* were grown with 0.4% glucose overnight, centrifuged at 4000 rpm for 4 min, washed twice with LB, and resuspended separately in LB with and without glucose with an initial OD_600_ of 0.1.

### 2.3 Sample preparation

Cells grown to mid-log phase were fluorescently labeled using 20 μM SYTO^®^ BC Green Fluorescent Nucleic Acid Stain (Thermo Fisher Scientific) in their growth medium, incubated at room temperature for at least 30 min without light exposure, and then centrifuged at 4000 rpm for 4 min. The cells were rinsed once and well mixed using a vortex mixer to remove excessive dye before being resuspended in the DEP buffer. The DEP buffer solution (final pH = 6.8) was prepared by adding deionized (DI) water to 1× phosphate-buffered saline until the solution conductivity was nearly 100 μS/cm. The DEP buffer also contains from 1 to 2% (v/v) glycerol to match the growth medium osmolarity.

### 2.4 Microchannel fabrication and preparation for electrokinetics

Microfluidic devices for electrokinetic mobility measurements and 3DiDEP analysis were fabricated in laser-cut poly(methyl methacrylate) (PMMA) chips using a CNC controlled micro-mill (Microlution), and bonded using a solvent-assisted bonding process (Wang et al., 2019) after cleaning with acetone, methanol, isopropanol and DI water in sequence. Both channel ends of the device were then bonded with electrical-insulating sleeve washers (McMaster-Carr) using a 5-minute two-part epoxy glue (Gorilla Glue). The sleeve washers serve as fluid reservoirs with a volume of ∼70 μL to contain bacterial samples. After bonding, each microchannel was connected to a syringe using Tygon tubing (McMaster-Carr) to flush and stabilize channel wall (“priming”) with the following reagents and conditions: 0.1 M potassium hydroxide, 1 mL/min, 10 mL; DI water, 1 mL/min, 10 mL; DEP buffer, 800 μL/min, 7 mL.

### 2.5 3DiDEP assay and image analysis

The geometry and dimensions of the 3DiDEP microchannel have been illustrated in Supplementary Figure 1. In brief, the microchannel has a cross-sectional area of 500 μm × 500 μm (width × depth) in its opening (non-constricting) region. At the center of the microchannel, the cross-sectional area is constricted to 50 μm × 50 μm (width × depth), resulting in a constriction ratio of one hundred. 3DiDEP trapping and derivation of trapping voltages have been described in previous studies (Braff et al., 2012, 2013; Wang et al., 2019). In brief, cells were resuspended in the DEP buffer at an OD600 ∼ 0.05 and quickly added into the 3DiDEP microchannel via the fluidic reservoirs. A “linear sweep” DC voltage difference increasing linearly with time at 1 V/s from 5 to 100 V was applied across the microchannel via an HVS-448 high-voltage power supply (LabSmith), controlled by a customized LabVIEW program. The SYTO^®^ BC fluorescence intensity increases with time as bacterial cells accumulate near the constricted region (Supplementary Video 1) and was recorded by time-lapse image sequences captured at 1 frame/s using a charge-coupled device (CCD) camera (CoolSNAP HQ2, Photometrics) fitted to an inverted fluorescence microscope (Eclipse Ti-U, Nikon). The fluorescence intensity data (background subtracted) near the 3DiDEP constriction versus time (i.e., the applied voltage) were fitted into a polyline with two segments, whose intersection point was taken as a variable optimized using the least squares method by a customized MATLAB R2019b (MathWorks) code. The applied voltage corresponding to the best-fit intersection point was extracted as the trapping voltage during 3DiDEP.

### 2.6 Linear electrokinetic mobility

To track the location of single cells and measure their linear electrokinetic mobility, a 500 μm × 60 μm × 1 cm (width × depth × length) straight microfluidic channel was designed and fabricated as described above. Prepared bacterial samples were loaded into the fluid reservoirs of a primed microchannel and fine-tuned until cell motion induced by gravity or pressure-driven flow was minimized. A pair of platinum electrodes (LabSmith) was connected to each reservoir of the channel ends to apply a constant DC voltage ranging from 10 to 50 V (with a 10 V increment) across the microchannel using an HVS-448 high-voltage power supply (LabSmith). During the linear electrokinetics, image sequences capturing cell movement in the microchannel was obtained using an inverted microscope (Eclipse Ti-U, Nikon) equipped with a sCMOS camera (Andor, Oxford Instruments) and image acquisition software (NIS-Element, Nikon). To minimize Joule heating effects, the DC voltage was applied for less than one min. To automatically extract cell locations and velocities in the obtained image sequences, a particle tracking algorithm was implemented using the python package trackpy (Allan et al., 2021) (Figure 1b). First, image sequences were loaded using a python package PIMS. Next, features of fluorescent single cells were identified and linked across the entire time frame, creating trace information for each detected cell (Crocker and Grier, 1996). Cell traces either appearing in less than five sequential frames or showing no increments in cell position with time were considered as noise traces and removed. K-means clustering was applied in noise filtering to exclude cells stuck to the channel wall. In order to derive linear electrokinetic mobilities and exclude background flow noise, we used a 3D fitting model to decouple nonlinear effects induced by pressure-driven or electrothermal flows. Data of single cell velocities, cell locations, and corresponding applied electric fields were interpreted based on a physical model with an equation,

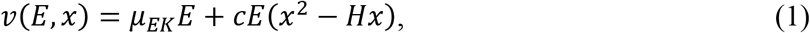

where *v* is the velocity, *Ε* is the magnitude of electric field, *x* is the distance of the cell measured from the bottom channel wall, *μ*_*EK*_ is the electrokinetic mobility, His the width of the microchannel, and *с* is a constant (Figure 1b and 1c). The quadratic term in the model equation represents a parabolic-shaped flow profile depending on the cell location *𝒳*, as an effort to decouple the background nonlinear flow effects (Wang et al., 2017). Using Equation 1, the best-fit parameter *μ*_*EK*_ was computed as the linear electrokinetic mobility for each strain.

### 2.7 Cell segmentation and polarizability

Cell cultures grown to mid-log phase were centrifuged at 4000 rpm for 4 min, and concentrated to OD600 ∼ 3. A SYTO^®^ BC Green Fluorescent Nucleic Acid Stain (Thermo Fisher Scientific) was added to the cell samples with a final concentration of 5 nM (0.1% v/v), and cells were stained in the dark at room temperature for 30 min. To immobilize the cells during imaging, 2 μL of the cell suspension was placed at the center of a 24 × 50 mm^2^ coverslip and covered by a thin 1% agarose pad (1 mm × 10 mm × 10 mm) (Skinner et al., 2013). Images were taken using the inverted microscope (Eclipse Ti-U, Nikon) with a 100X or 150X objective.

With the obtained images, cell segmentation was performed using Coli-Inspector (Vischer et al., 2015), an add-on project to an image analysis software (ImageJ) and its plugin (ObjectJ). At least 400 individual cells were identified to measure their length and diameter (i.e., long and short axis). The measurements were used to calculate the Perrin friction factor (*ξ*) and cell shape factor (Wang et al., 2019). Average cell shape factor and DEP mobility for each strain were used to derive the Clausius-Mossotti factor, a measure of the cell polarizability (see Supplementary Figure 2 and Supplementary Note 3.2).

### 2.8 Colistin susceptibility and growth

*E. coli* strains varying in LPS composition were investigated for susceptibility to colistin. Each strain was grown overnight (see section “Bacterial strains” in Methods) and diluted to OD600 ∼0.1. To prepare a minimum inhibitory concentration (MIC) assay, colistin was added to 150 μL LB containing streptomycin (replaced by spectinomycin for the O-antigen expressing strain) per strain to reach a final concentration ranging from 0.0156 to 2 μg/mL (with a two-fold increment) in a clear flat-bottom 96-well plate (Corning). Then, 1.5 μL aliquots of bacterial samples were added to the multiwell plate and covered with a sealing film (Breathe-Easy). Cell abundance in the multiwell was measured every 10 min by OD_600_ using a microplate spectrophotometer (Epoch 2, Biotek). Each microplate was incubated at 37°C for 20 h in shaking mode. Three biological samples were repeated for each strain and colistin concentration. Based on the bacterial abundance at the final time measurement, MIC of colistin was determined by strain turbidity, and cell growth was analyzed as a function of *𝒳*μg/mL colistin concentration with the following equation,

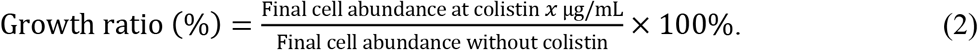

A maximal growth rate was calculated based on the time-series abundance. First, local growth rates were computed as the time derivatives of the natural logs of cell OD_600_ using data points collected every three timepoints. Then the maximum local growth rate calculated within the entire 20 h time window was obtained for each studied strain.

### 2.9 Outer membrane permeability assay

An *N*-phenyl-1-naphthylamine (NPN) uptake assay was performed to compare the effects of differential LPS structures on colistin-mediated permeabilization of outer membranes in the studied strains using a protocol adapted from the previous study (MacNair et al., 2018). In brief, bacterial cells in mid-log phase were washed twice with 5 mM HEPES buffer (pH ∼7.2, Sigma-Aldrich) containing 20 mM glucose and adjusted to have their OD600 reaching about 0.4. One hundred microliters of cells were added to 100 μL HEPES buffer containing 20 μM NPN and a varying concentration of colistin (ranging from 0.39 to 200 μg/mL, two-fold of the final concentrations) in a black clear flat-bottomed 96-well plate. After allowing the microplate to sit for an hour at room temperature without light exposure, NPN fluorescence levels were read using the Varioskan Flash plate reader (Thermo Scientific) with an excitation and emission wavelengths of 355 ± 5 nm and 420 ± 5 nm, respectively. Three biological replicates were measured for each strain and colistin concentration. The NPN uptake factor was calculated as the ratio of background-subtracted fluorescence (subtracted by the fluorescence levels in the absence of NPN) values of the bacterial suspension and of the suspending medium (Helander and Mattila-Sandholm, 2000). Then, the NPN uptake factor was normalized by a cell abundance factor to exclude the effect of fluorescence increase induced by cell growth. The cell abundance factor was estimated using the maximal growth rate derived from the colistin susceptibility assay as described above (see section “Colistin susceptibility and growth” in Methods). The cell abundance factor is larger than 1 for cases exhibiting a positive growth rate where colistin concentration was lower than the MIC. Finally, fractional changes of normalized NPN uptake (Figure 5) for the O-antigen expressing strain and each core mutant strain relative to the parental strain were calculated as

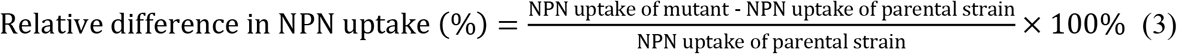

to analyze the effects of LPS structural variations on outer membrane permeability to colistin.

### 2.10 Statistical analysis

Unless otherwise mentioned, a Kruskal-Wallis test was used to compute the statistics across all the studied strains as the input of a multiple comparison test where a Tukey’s honestly significant difference criterion was applied to determine which groups are significantly different in linear electrokinetic mobility, trapping voltage, DEP mobility, polarizability, and MIC for each LPS structural condition. To compare timewise growth measurements between the studied strains, a two-way analysis of variance (ANOVA) was performed to assess how LPS structural variation and colistin concentration affect cell abundance. At least three biological replicates were used for all the statistical tests. An R package dplyr v1.0.7 (Wickham et al., 2022) and R function (aov) were used to perform the two-way ANOVA analysis, and MATLAB 2019b was used to perform the multiple comparison and Kruskal-Wallis test.

## 3 Results and Discussion

### 3.1 LPS compositional variations drive changes in *E. coli* electrokinetic properties

Due to the presence of LPS and other external appendages, bacteria moving in electrolyte solutions act as so-called soft particles, i.e., hard particles covered by an ion-permeable polyelectrolyte layer (PEL) (Duval and Ohshima, 2006; Ashrafizadeh et al., 2020), in which both the distribution of charge density and the arrangement of LPS monomers can substantially affect cell electrokinetic properties (Ohshima, 2013). Not only the spatial details of phosphate groups in lipid A and LPS inner core but also electroneutral sugar residues in the outer core and O-antigen (Figure 1a) confer an inhomogeneous distribution of polyelectrolyte segments within the PEL. According to the electrokinetic theory of diffuse soft particles by Duval and Ohshima (Duval and Ohshima, 2006; Ohshima, 2013), electrophoretic mobility of a soft particle increases with an increasing volumetric charge density (*ρ*_*PEL*_) or a decreasing δ, where δ represents the degree of inhomogeneity measured by the width of distribution of uncharged segments near the front edge of the PEL. In addition, PEL segments can be considered as resistance centers exerting Stokes’ drags on the liquid flowing within the PEL with a frictional coefficient *γ ∼ η*/(1/*λ*)^2^, η being the viscosity of the electrolyte solution (Duval and Ohshima, 2006; Ohshima, 2013; Ashrafizadeh et al., 2020). The electrophoretic softness, 1/λ, is a characteristic penetration length representing how easily the fluid can flow through the PEL. Especially for the low electrolyte concentration used in this study, electrokinetic behaviors of PEL become increasingly sensitive to the spatial distribution of the polyelectrolyte segments (Duval and Ohshima, 2006).

We hypothesize that LPS structural differences in bacteria can be manifested in their PEL volumetric charge density (*ρ*_*PEL*_), inhomogeneous distribution of polymer segments (δ), and electrophoretic softness (1/λ), thereby leading to distinguishable electrokinetic behaviors. To test this hypothesis, we studied several *E. coli* K-12 strains with different LPS structures. To model scenarios with reduced PEL thickness and altered charge conditions, we studied three “deep-rough” LPS mutants (Δ*waaC*, Δ*waaF*, and Δ*waaP*) (Halvorsen et al., 2021) that are unable to synthesize the full LPS core (Figure 1a). Previously, it has been reported that a deletion of *waaC* results in a loss of the majority of the LPS core including the inner core phosphates (Gronow et al., 2000). WaaF transfers a heptose II to the inner core (Gronow et al., 2000), and its deletion causes a severely truncated LPS core but maintains heptose I phosphates in the inner core. WaaP deletion mutants possess a nearly intact oligosaccharide core, but lack heptose III and inner core phosphate groups (Yethon et al., 1998; Raetz and Whitfield, 2002) (Figure 1a). These expected structural changes in core oligosaccharide have been confirmed for each studied strain (Supplementary Table 1) previously by total LPS extraction and SDS-PAGE (Halvorsen et al., 2021). In addition to a wild type control (parental strain in Figure 1), we included a strain with additional O-antigen synthesis to model PEL with decreased charge density and enhanced polymer segment inhomogeneity (Feldman et al., 1999). The presence of O-antigen in this strain has also been confirmed in the prior study (Halvorsen et al., 2021).

To detect LPS structural variations by whole-cell electrokinetic mobility, we employed a single-cell tracking algorithm (see Methods) to extract the dependence of cell migration on the applied electric field. The algorithm captures velocity, position within the microchannel, and the corresponding applied voltages for each individual cell, and fits them using a 3D regression model to decouple the confounding effects of pressure-driven flow and nonlinear electrokinetic flow (Figure 1b and 1c). Cell motion in the straight microchannel is driven by both the background electroosmotic flow and electrophoresis of the cell. However, the background electroosmotic flow is constant given the same counterion concentration and applied electric field at the microchannel/medium interface (Kirby and Hasselbrink, 2004), leaving the linear electrokinetic mobility purely a function of cell electrophoresis. Note that the cell tracking analysis and the 3DiDEP cell trapping analysis in the following section require cell staining with a cationic fluorescent nucleic acid dye (SYTO® BC) (Thermo Fisher Scientific Inc., 2014). While an excessive amount of the SYTO stain can impact the cell membrane composition and morphology (Deng et al., 2020), we used a dye concentration per cell ∼10^4^ times lower than the value expected to induce a noticeable membrane alteration (see Supplementary Note 3.3 for details).

As shown in Figure 1d, the linear electrokinetic mobility of all the tested strains is negative, which agrees with the electric properties of the charged groups (e.g., carboxyl and phosphate groups) in the cell outer membrane and the LPS structures (Adams et al., 2014; Liang et al., 2016). O-antigen restoration results in a significantly less negative (*p* = 0.00024) electrokinetic mobility compared to the control strain lacking O-antigen. This observation is consistent with our hypothesis that the longer electroneutral oligosaccharide on this strain may decrease electrokinetic mobility by not only increasing the PEL thickness, thereby lowering the volume charge density, but also by enhancing the inhomogeneity of the polyelectrolyte segment distribution in the PEL.

In contrast, the truncated LPS core present in the Δ*waaF* and Δ*waaP* mutants enhances cell electrokinetic mobility relative to the parental strain by 48.7% (*p* = 0.0023) and 22.7% (*p* = 0.012), respectively (Figure 1d). This is consistent with the increased electrophoretic mobility of bacteria due to LPS removal after an EDTA treatment observed previously (Chandraprabha et al., 2009). These results indicate that truncation of LPS core components can lead to an increased PEL charge density, and thus a higher electrokinetic mobility. For instance, the truncated core of Δ*waaF* mutants contains an additional phosphate group compared to Δ*waaP* cells (Figure 1a), which is predicted to increase charge density and result in a lower inhomogeneity of polyelectrolyte segment distribution in the PEL relative to the control strain. Correspondingly, the Δ*waaF* mutant exhibits the highest electrokinetic mobility among tested strains (Figure 1d). In contrast, Δ*waaP* mutants are predicted to have the lowest charge density relative to the control due to the loss of both phosphates and heptose III residues.

The Δ*waaC* mutation results in a more drastic LPS truncation (Figure 1a), and correspondingly shows a significant decrease in electrokinetic mobility relative to the Δ*waaF* and Δ*waaP* strains (*p* = 0.007 and 0.003, respectively). Though it contains the most extensive LPS truncation, the Δ*waaC* mutant exhibits an electrokinetic mobility comparable to that of the parental strain with an intact core (*p* = 0.45). We hypothesize that a *waaC*-deficiency generates an increased charge density but a decreased electrophoretic softness (1/λ), resulting in stronger frictional forces exerted on the liquid flowing within the PEL, which could be compensatory to each other in terms of their effects on the electrokinetic mobility. Comparable electrophoretic mobilities have been reported previously between *E. coli* constitutively producing long (10 – 100 μm) F pili and Ag43 proteins (Francius et al., 2011), suggesting that cell electrokinetic mobility is affected both by the length of cell surface appendages and the interplay between electrophoretic softness and spatial arrangement of charged groups of the PEL. In deep-rough LPS mutants, physiological alterations may also contribute to strain-specific signatures that are differentiable by electrokinetic mobility measurement. For instance, Δ*waaC* and Δ*waaF* mutations have lower levels of OmpC porin in the outer membrane, whereas the Δ*waaP* mutant produces wild-type levels of OmpC (Nakao et al., 2012; Halvorsen et al., 2021). The extracellular loops of OmpC are negatively charged to promote the porin’s cation transport function (Kojima and Nikaido, 2014). Thus, the lower abundance of OmpC in Δ*waaC* and Δ*waaF* cell outer membrane may have partially compensated for the effect of LPS core truncation on charge density of surface soft layers, thereby lowering their baseline values of electrokinetic mobility. It also reveals that characterizing LPS properties on whole cells is beneficial for capturing physiological changes that are difficult to observe when studying LPS alteration using purified LPS alone.

### 3.2 Trapping *E. coli* based on LPS structures using 3DiDEP

In addition to studying cell electrokinetic mobility, a property induced by cell surface charge, we next investigated cell polarizability, which is the tendency of electric dipoles to form at the cell surface (not necessarily charged) when subjected to externally applied electric fields. Cell polarizability has been correlated with the extracellular electron transfer capability in electrochemically active bacteria in our previous study (Wang et al., 2019), and has been proven to be a sensitive label-free biomarker to detect variations in cell outer membrane components (Braff et al., 2013).

To measure cell polarizability, we used a microfluidic 3DiDEP trapping technique, incorporating linear sweep analysis, in which the applied electric field increases linearly with time across the microchannel (Figure 2a). The microchannel features a 3D hourglass shaped constriction at the middle, which bridges two main channels (Supplementary Figure 1). The 3D constriction has a cross-sectional area 100 times smaller than that of the main channel, which creates a strong electric field gradient to enable a high DEP force for immobilizing bacteria near the constriction (Hawkins et al., 2007; Pysher and Hayes, 2007; Moncada-Hernandez et al., 2011; Jones et al., 2015; Crowther et al., 2019). In response to the linear sweep voltage, bacteria flowing down the main channel start to slow down and gradually accumulate near the constriction when DEP balances cell motion due to linear electrokinetics along the electric field direction (Supplementary Video 1). The threshold voltage for the onset of cell immobilization, termed the ‘trapping voltage’, is a measure of the comparison between cell DEP mobility and the counteracting linear electrokinetic mobility. The criterion for 3DiDEP immobilization of a single cell can be expressed as (Moncada-Hernandez et al., 2011; Braff et al., 2012; Jones et al., 2015; Lapizco-Encinas, 2019; Wang et al., 2019)

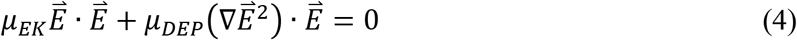

where *μ*_*EK*_ is the linear electrokinetic mobility, 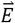 is the local electric field corresponding to the measured 3DiDEP trapping voltage, and *μ*_*EX*_ is the DEP mobility. The studied *E. coli* K-12 strains are rod shaped and can be modeled as ellipsoidal particles with semi-axes *a* > *b* = *c* (Khoshmanesh et al., 2012) (Supplementary Figure 2). Thus, the DEP mobility *μ*_*EX*_ can be expressed as

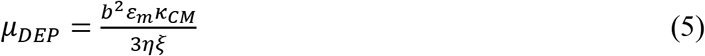

where *ε*_*m*_ and *η* are the permittivity and viscosity of the surrounding medium, respectively, and *ξ* is the Perrin friction factor (see Supplementary Material 3.2 for a detailed derivation). The Clausius-Mossotti factor, *κ*_*CM*_, is a measure of the relative polarizability of the cell compared to the surrounding medium, and can be estimated from the experimentally determined ‘trapping voltage’ required for initiating 3DiDEP cell trapping.

**Figure 2.**
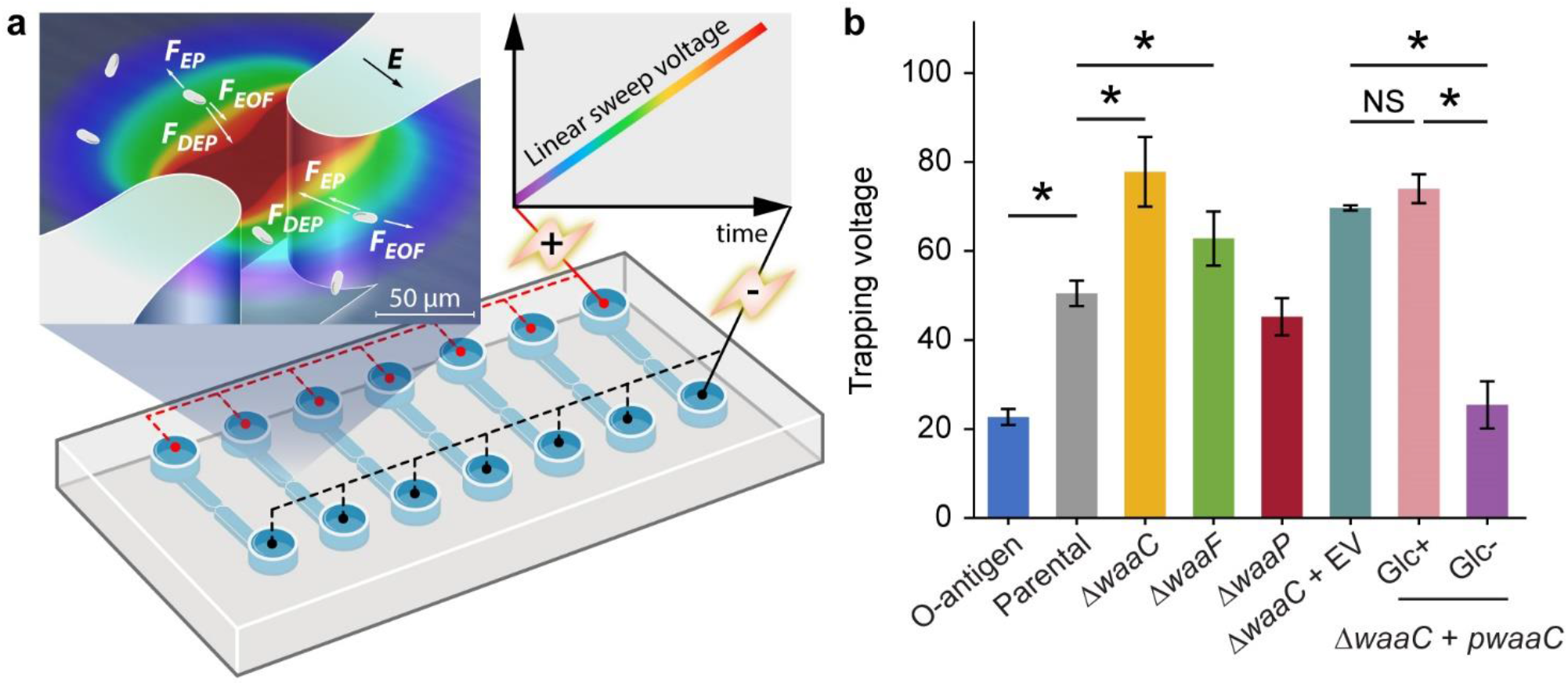
Immobilizing LPS mutants using 3D insulator-based dielectrophoresis (3DiDEP). (a) Schematic of a microfluidic device with an array of 3DiDEP microchannels. Dimensions of the 3D microchannel geometry are shown in Supplementary Figure 1. A DC potential difference increasing linearly with time at 1 V/s was applied across the channel. Magnified view of the 3D constricted region depicts the 3DiDEP trapping principle: bacteria near the constriction are immobilized when the DEP force balances drag forces due to background electroosmotic (EOF) flow and electrophoresis (EP). Electric field intensity is illustrated in the background color scale (dark red indicates higher values). (b) Measured trapping voltage (threshold voltage at the onset of cell trapping) for each strain. Error bars indicate standard deviations. Asterisks indicate statistical difference (\**p* < 0.05, NS: not significant).

Several prior studies on insulator-based electrokinetic systems reported significant effects of nonlinear electrophoresis on particle/cell motion due to the application of strong electric fields (Tottori et al., 2019; Antunez-Vela et al., 2020; Cardenas-Benitez et al., 2020). While these studies demonstrate good agreement between particle velocity measurements and a nonlinear electrophoresis model (Schnitzer and Yariv, 2012; Schnitzer et al., 2013), cell trapping in our 3DiDEP system presents a distinctive physical scenario for several reasons. First, given the low zeta potential of the PMMA microchannel and high surface charge density of the examined bacterial cells, electrophoresis exceeds electroosmosis in our system, which means that cell trapping is impossible without the contribution of DEP. Second, the DEP force created at a 3DiDEP constriction is over 100 times stronger than the force generated in a corresponding 2D insulating constriction (Supplementary Figure 3), avoiding the use of large correction factors that is often required in 2DiDEP systems (Hill and Lapizco-Encinas, 2019; Cardenas-Benitez et al, 2020). Third, using a combination of the SY model for nonlinear electrophoresis (Schnitzer & Yariv, 2014) and COMSOL simulations of electric field distributions solved for each studied strain based on their measured trapping voltages, we found that the nonlinear component of electrophoretic velocity is an order of magnitude smaller than its linear counterpart for each strain investigated (Supplementary Table 3 and Supplementary Figure 4). Taken together, we believe we can neglect the effect of nonlinear electrophoresis in our 3DiDEP cell trapping analysis. Modeling and evaluation of potential effects of nonlinear electrophoresis on the cell motion is discussed in Supplementary Note 3.1.

In addition, we show that the strong electric field generated at the 3DiDEP constriction does not significantly impact cell membrane integrity and cell viability. First, our simulation results using a previously developed 3D time-dependent COMSOL model (Wang et al., 2017) show that the maximum temperature rise induced in the 3DiDEP channel is less than 3 °C for a trapping voltage lower than 100 V, suggesting small Joule heating effects (Supplementary Figure 5). In addition, using both of a previously developed microfluidic screening assay (Garcia et al., 2016; Garcia et al., 2017) and commercially available electroporation cuvettes, we found that the threshold electric field (∼10 kV/cm) to induce a transmembrane potential that is adequate to permeabilize the cell membrane is 1 – 2 folds higher than the maximum electric field generated in our 3DiDEP experiments (see Supplementary Figure 6 and Supplementary Note 3.4 for details). These findings highlight the potential of 3DiDEP as a noninvasive method for characterizing surface properties (e.g. LPS) of bacterial cells.

Our 3DiDEP measurement reports an inverse relationship between the length of *E. coli* K-12 LPS and the trapping voltage for 3DiDEP cell immobilization (Figure 2b). The O-antigen expressing strain requires a 55% lower trapping voltage relative to the parental strain (*p* = 0.025), and correspondingly this strain expresses longer LPS with distal O-antigen repeats. In addition to the lower electrokinetic mobility resulting from the O-antigen expression (Figure 1d), this can be explained by a higher polarizability thereby a lower trapping voltage required to counteract with linear electrokinetics, which will be shown in the following section. In contrast, the Δ*waaC* mutant strain requires the highest trapping voltage for 3DiDEP immobilization (*p* = 0.025, Figure 2b). To confirm this relationship, we then measured trapping voltages of a Δ*waaC* mutant strain complemented with an empty vector (Δ*waaC* + EV) and a *waaC* mutant strain complemented with pCH450-*waaC* grown with (Δ*waaC* + *pwaaC*, Glc+) or without glucose (Δ*waaC* + *pwaaC*, Glc-). Removal of glucose in the growth medium induces *waaC* expression and restores synthesis of the wild-type core (Figure 1a). As expected, when grown with glucose (Glc+ in Figure 2b), the Δ*waaC* + *pwaaC* strain shows a comparable trapping voltage (*p* = 0.077) relative to the isogenic control (Δ*waaC* + EV). Removal of glucose (Glc-) in the growth medium restores the full-length LPS core, which results in a 65.7% drop in the trapping voltage (*p* < 0.05), further indicating that cells lacking *waaC* require a stronger trapping voltage for 3DiDEP immobilization. The Δ*waaC* + EV strain (*p* = 0.18) and the Δ*waaC* + *pwaaC* strain grown with glucose (*p* = 0.65) show comparable trapping voltage relative to the parental strain, suggesting that the introduction of the vector pCH450 does not significantly affect the cell polarizability. Furthermore, deletion of *waaF* and *waaP* lead to a 24.3% increase (*p* = 0.025) and a 10.5% decrease (*p* = 0.053) in trapping voltage relative to the parental strain, respectively. Although the Δ*waaF* mutation should result in only one fewer phosphorylated heptose compared to Δ*waaC* cells, the Δ*waaF* mutant exhibits a trapping voltage ∼20% lower than that of the Δ*waaC* mutant. Comparison between the trapping voltages of Δ*waaC* and Δ*waaF* strains suggests that 3DiDEP trapping is sensitive enough to detect a single heptosylation event. These data indicate that 3DiDEP can be used to selectively immobilize bacteria based on their LPS length.

### 3.3 LPS structural variations affect cell DEP mobility and polarizability

To determine the cell polarizability, for each trapping voltage measurement, we performed additional particle image velocimetry (PIV) analysis (Wang et al., 2019) for cell motion in the corresponding main channel region where the electric field is uniform. Electrokinetic mobilities measured by this PIV analysis show a high consistency with the results measured using our cell tracking algorithm (Supplementary Figure 7). Incorporating the electrokinetic mobility and the corresponding threshold electric field numerically computed from the measured trapping voltages (Equation 4), we derived the DEP mobilities for all the tested strains (Figure 3a). To decouple the effects of cell size/shape, we computed the Clausius-Mossotti factor (*κ*_*CM*_) as a measure of cell polarizability (Equation 5, Figure 3b) using cell shape parameters (*b* and *ξ*) measured for all the tested strains using high magnification microscope images (see Methods).

**Figure 3.**
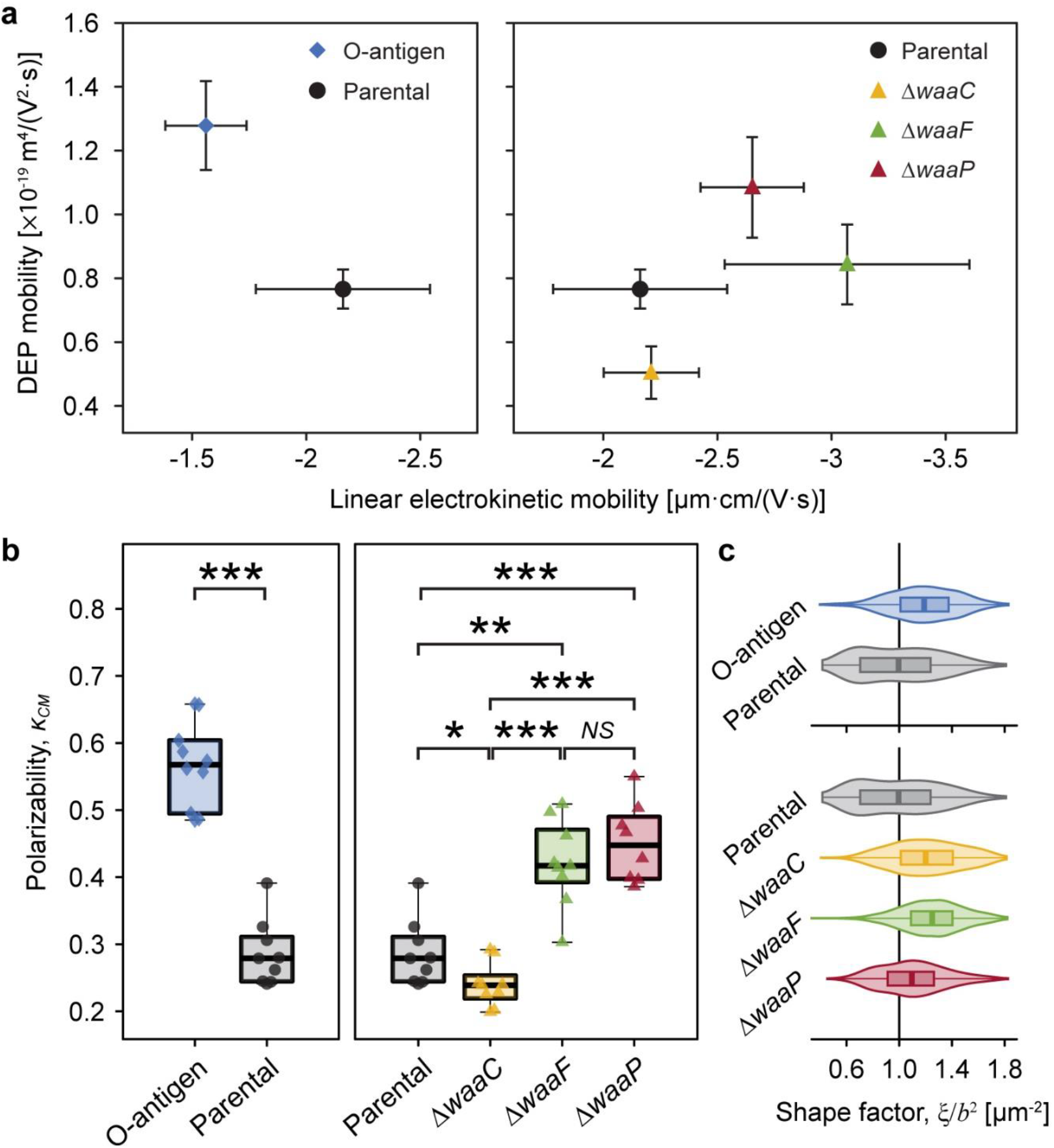
Strong correlation between *E. coli* LPS structural variations, DEP mobilities, and polarizability. (a) Plot of measured DEP mobilities against linear electrokinetic mobilities for each strain indicates that *E. coli* expressing O-antigen (diamond, left) and core synthesis mutants (triangles, right) clearly deviate from the parental strain (circle). Error bars indicate standard deviations. Statistical difference of DEP mobilities is shown in Supplementary Figure 8. (b) Cell polarizability, as measured by the Clausius-Mossotti factor (*κCM*) of each strain. Asterisks indicate statistical difference (\**p* < 0.02, \*\**p* < 0.005, and \*\*\**p* < 0.001, NS: not significant). (c) Cell shape factor, *ξ*/*b*^2^, of studied strains, where *ξ* is the Perrin friction factor (Koenig et al., 1975) and *b* is the short semi-axis of the cell. The shape factor represents how much the ellipsoidal cell shape influences the drag force versus the DEP force. See Supplementary Note 3.2 for a detailed explanation.

Our results show that the O-antigen expressing strain possesses a higher DEP mobility (by 66.8%, *p* = 0.025, Figure 3a and Supplementary Figure 8) and a higher *κ*_*CM*_ (by 98.1%, *p* = 0.00024, Figure 3b) but a lower electrokinetic mobility (by 27.8%, *p* = 0.00024) relative to the parental strain (Figure 1d), suggesting that the presence of O-antigen not only lowers the negative charge density but also enhances the polarizability of the cell surface. In contrast, although knocking out *waaC* does not significantly affect cell electrokinetic mobility (Figure 1d), its removal decreases the DEP mobility by 34.2% (*p* = 0.025) and polarizability by 16.4% (*p* = 0.015). As discussed above, the presence of LPS adds a charged soft PEL layer around a bacterium. Ion transport in this soft PEL layer and electrical double layer jointly contribute to the induced dipole moment coefficient. As demonstrated in our previous study (Dingari and Buie, 2014), fibrillated strains with a thicker PEL have a higher induced dipole moment coefficient and thereby are more polarizable than unfibrillated strains in the low electrolyte concentration regime, which agrees with our observation that long LPS chains resulting from O-antigen expression leads to a higher polarizability but truncating most of the LPS core via *waaC* deletion results in a significantly lower polarizability (Figure 3b). Interestingly, deletion of *waaP* increases all three parameters that we tested, showing an enhanced DEP mobility (by 41.6%, *p* = 0.025), polarizability (by 57.6%, *p* = 0.00076), and electrokinetic mobility (by 22.7%, *p* = 0.012). By comparison, the mutant strain lacking *waaF* exhibits the strongest increase in cell electrokinetic mobility (*p* = 0.0023) among all of the examined LPS mutations, a 47.3% increase in polarizability (*p* = 0.0013), but small changes in DEP mobility (*p* = 0.18). These results indicate that not only the PEL thickness but also the polyelectrolyte segment distribution drive changes in cell polarizability. The polar to nonpolar composition of the LPS chains affects the conductivity distribution in the PEL, thereby influencing cell polarizability, which is consistent with the discrepancies observed between the parental strain and mutants lacking *waaF* and *waaP* (Figure 3b). It is interesting to note that the three core mutants exhibit altered morphology relative to the parental strain (Figure 3c). Because cell shape can affect both the drag force via the Perrin friction factor (*ξ*) (Koenig, 1975) and the DEP force via the short semi-axis (*b*) of the cell, the ratio *ξ*/*b*^2^ (Figure 3c) represents how much the ellipsoidal cell shape influences the drag force versus the DEP force (Wang et al., 2019). Note that deletion of *waaF* does not affect DEP mobility significantly but drastically increases polarizability, indicating that the enhanced polarizability can also be attributed to the morphological change induced by this mutant.

### 3.4 Outer membrane permeability increases with cell polarizability

After characterizing the differential electric and dielectric properties of *E. coli* strains conferred by their LPS structural variations, we next sought to examine whether changes in polarizability and electrokinetic mobility are associated with variations of outer membrane permeability against membrane-disrupting reagents, such as antibiotics. We hypothesize that cells capped with thick PEL layers with small electrophoretic softness, i.e., less penetrable by liquid flow, carry less permeable outer membranes, and thereby are distinguishable by an increased cell polarizability and altered electrokinetic properties. To test this hypothesis, colistin was chosen as the antibiotic agent, as it is known to interact with the bacterial outer membrane by destabilizing LPS (Sabnis et al., 2021). The activity of colistin starts with electrostatic interaction between the positive charges carried by colistin and the negatively charged phosphates in LPS, which is believed to displace divalent cations (Ca^2+^ and Mg^2+^) that are important to bridge and stabilize the LPS layer (Velkov et al., 2010).

To determine cell susceptibility to colistin, we measured minimum inhibitory concentrations (MIC) and cell growth (cell abundance normalized by values when colistin is absent) across a varying colistin level for each strain (Figure 4). As expected, *E. coli* expressing O-antigen shows the highest MIC (Figure 4a) and the highest growth ratio at all colistin levels (Figure 4b) among the tested strains. The parental strain shows an MIC of 0.33 μg/mL and a cell abundance decreased by 80% when the colistin concentration reaches 0.25 μg/mL, whereas the O-antigen expressing strain shows a two-fold increase in MIC compared to the parental strain, and keeps a high abundance (99%) at 0.25 μg/mL colistin. The enhanced resistance to colistin suggests that O-antigen facilitates the barrier function of the outer membrane against colistin, which is consistent with the high polarizability and low electrokinetic mobility observed for this strain. Electrostatic interactions between the cationic groups in colistin and phosphate groups in lipid A play important roles in peptide binding (MacNair et al., 2018). The presence of O-antigen decreases cell electrokinetic mobility (Figure 1d) and increases polarizability (Figure 3b), which indicates a lower charge density and/or a higher friction factor in the soft PEL layer, and thereby may have shielded the electrostatic effects and impeded colistin activity. In contrast, mutants lacking genes *waaC, waaF* or *waaP* exhibited a significantly lower growth in response to colistin compared to the parental strain (*p* = 4 × 10^−7^, 3.8 × 10^−7^, and 0.0006 respectively, two-way ANOVA), which can be explained by the absence of outer core oligosaccharides. Among these deep-rough LPS mutants, the Δ*waaC* strain shows the lowest MIC followed by the Δ*waaF* and Δ*waaP* strains (Figure 4a, *p* = 0.0005). A similar trend was found in colistin-mediated cell lysis of the Δ*waaC, ΔwaaF* and Δ*waaP* strains (Figure 4b) as well as in their DEP mobilities and polarizabilities (Figure 3a and 3b). These results suggest a correlation between cell susceptibility to colistin and LPS composition, which is distinguishable by their polarizability and electrokinetic properties.

**Figure 4.**
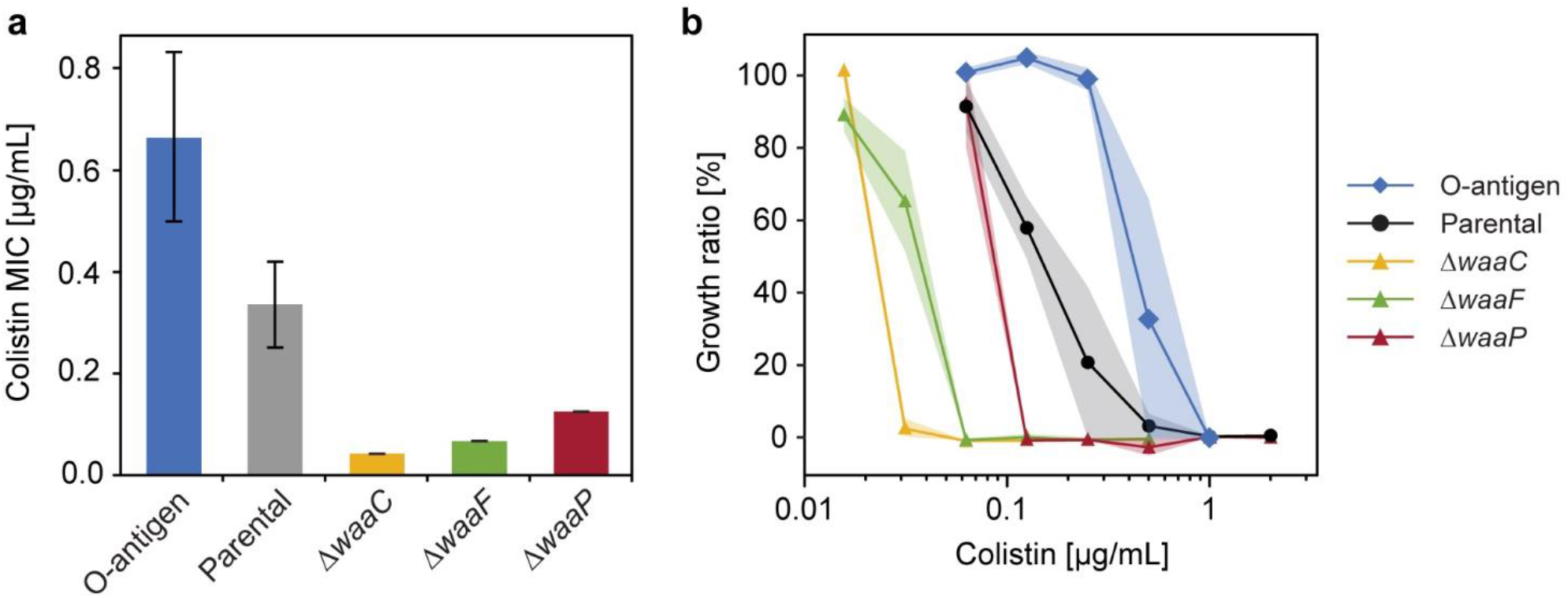
Variations in *E. coli* LPS structures affect bacterial response to colistin. (a) Minimum inhibitory concentration (MIC) of colistin tested for *E. coli* strains expressing different LPS structures (*n* = 3, error bars indicate standard error). (b) Growth inhibition of *E. coli* O-antigen expressing strain (diamonds), parental strain (circles), and mutants with truncated core oligosaccharide (triangles) exposed to varying colistin concentrations. Shaded areas indicate standard errors of three replicates.

To further investigate the correlation between cell electrokinetic properties and outer membrane permeability, we measured the uptake of *N*-phenyl-1-naphthylamine (NPN), a nonpolar probe that fluoresces in the hydrophobic environment of lipid bilayers (Helander and Mattila-Sandholm, 2000), for each strain in the presence of colistin (Figure 5). An intact outer membrane prevents NPN from entering the phospholipid layer and the subsequent fluorescence (MacNair et al., 2018). To examine if cells that differ in polarizability and electrokinetic mobility can be distinguished by their differential outer membrane permeability, we treated the *E. coli* strains with NPN and various concentrations of colistin, and measured the fluorescence after one hour of incubation. To decouple the possible effects of increased cell abundance (due to low colistin concentrations, Figure 4b) on NPN fluorescing, NPN uptake values were normalized by a cell abundance factor estimated from the maximal growth rate (Supplementary Figure 9, see Methods).

Figure 5a and 5b show the relative differences of NPN uptake calculated for each tested strain compared to the parental strain. As expected, the O-antigen expressing strain shows a strong decrease in its NPN uptake, by a factor of 15.7% to 53.3% compared to the parental strain, across all colistin concentrations (Figure 5a), indicating that the presence of O-antigen prevents NPN from entering the outer membrane. Conversely, mutants harboring truncated LPS show an increased NPN uptake even in the absence of colistin, indicating a compromised outer membrane with enhanced permeability to NPN. In addition, it has been reported that mutants expressing severely truncated LPS cores fail to insert many outer membrane proteins and consequently have to fill the “void” in the outer membrane with phospholipids (Nikaido Hiroshi, 2003), which also explains why their NPN fluorescence is stronger than the control even without colistin exposure (Figure 5b). Moreover, we observed a lower tolerance for colistin-mediated membrane disruption in the *waa* mutants compared to the parental strain, and the membrane disruption dynamics vary at different colistin levels. For instance, at a colistin concentration of 0.195 μg/mL, a stronger increase of NPN uptake was observed by roughly two-fold for Δ*waaC* and Δ*waaF*, and 67.5% for Δ*waaP*, respectively. The enhancement of NPN uptake caused by these mutants gradually decreases when colistin concentration increases up to 1 μg/mL (Figure 5b). This agrees with the distinct colistin susceptibility between the parental and mutant strains shown in Figure 4b. For example, cells are completely depleted at 0.031, 0.063, and 0.125 μg/mL colistin for Δ*waaC, ΔwaaF*, and Δ*waaP*, respectively, suggesting rapid cell lysis. However, cell depletion of the parental strain initiates at 0.063 μg/mL colistin and a complete cell clearance occurs at a concentration higher than 1 μg/mL (Figure 4b). As a result, at a low colistin level (0.195 μg/mL in Figure 5b), the three core mutants are hypersensitive to colistin and are immediately lysed, whereas the parental strain keeps an intact outer membrane to resist NPN, leading to the strongest NPN uptake discrepancy between mutants and the parental strain among the tested colistin levels (Figure 5b). At an intermediate colistin level (0.195 to 0.781 μg/mL in Figure 5b), because the rate at which cell lysis occurs for the parental strain increases (Figure 4b) while the mutants remain hypersensitive to colistin, the NPN uptake discrepancy between the mutants and the parental strain decreases with an increasing colistin concentration. When colistin exceeds 1 μg/mL, discrepancies between the NPN uptake for the mutants (Δ*waaC, ΔwaaF*, and Δ*waaP*) and the parental strain are small (Figure 5b), indicating that both parental and mutant strains are lysed quickly at this colistin level, and consequently exposing comparable amounts of phospholipids to NPN. The stronger increase of NPN uptake observed in Δ*waaC* and Δ*waaF* but not Δ*waaP* is consistent with the amount of intact core residues in each mutant (Figure 1a). Figure 5c compares the changes of cell electrokinetic properties and outer membrane permeability induced by LPS structural variations. It suggests that increased outer membrane permeability has opposite effects on cell linear electrokinetic mobility and polarizability. In addition, the trend of 3DiDEP trapping voltage matches the trend of NPN uptake, indicating the potential of using 3DiDEP to distinguish cells based on their outer membrane permeability. Together, *E. coli* outer membrane permeability and small molecule uptake strongly correlate with LPS compositional variations, thereby are distinguishable by cell polarizability and electrokinetic mobility (Figure 5c).

**Figure 5.**
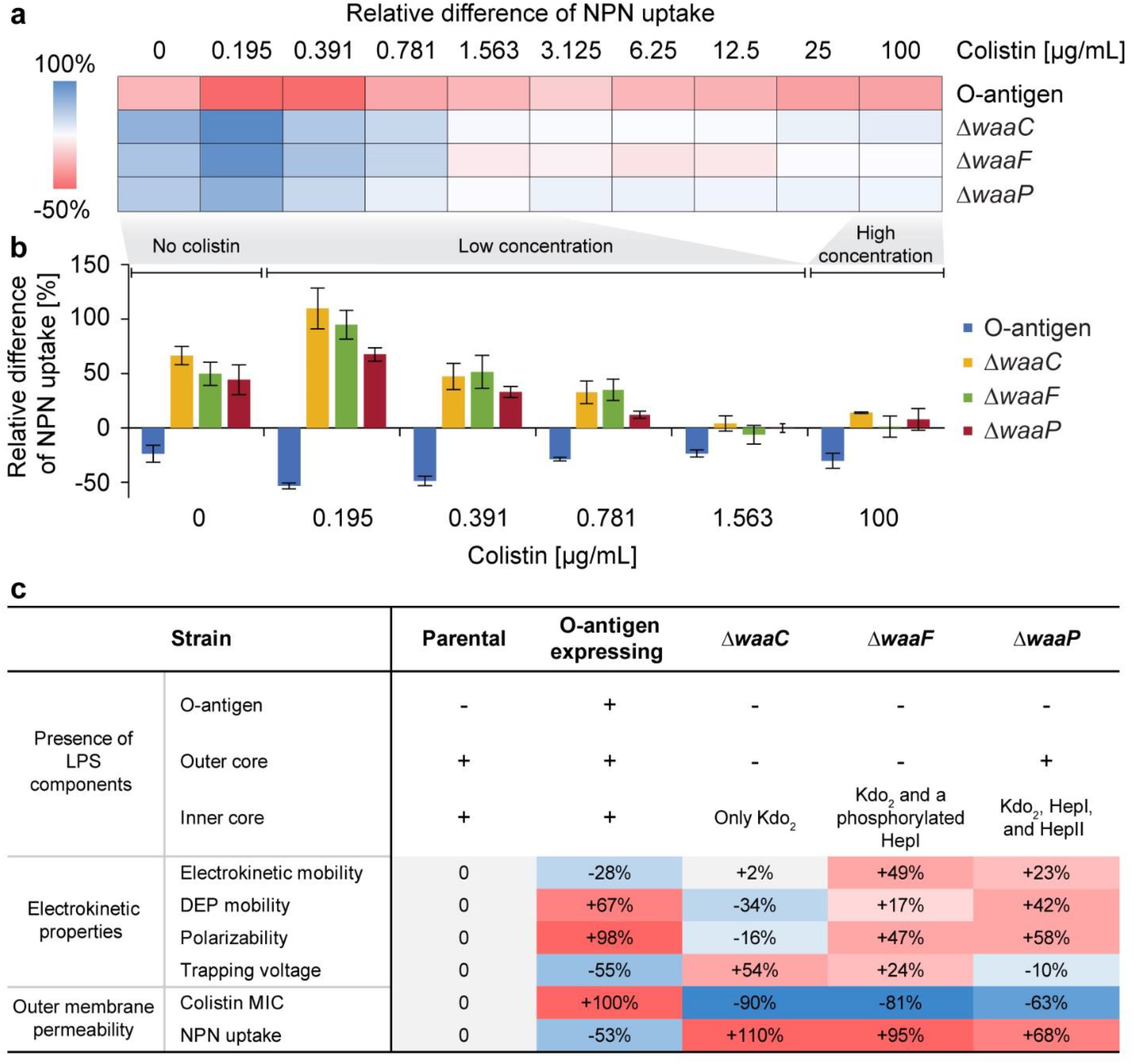
Effects of *E. coli* LPS structural variations on colistin-mediated outer-membrane disruption. (a) A heat map showing the mean relative differences of *N*-phenyl-1-naphthylamine (NPN) uptake compared to the parental strain for the *E. coli* O-antigen expressing strain and mutants with truncated core oligosaccharide (as computed by Equation 3 in Methods). NPN uptake represents the ratio of background (absence of NPN) subtracted fluorescence values of the bacterial suspension to that of the suspending medium normalized by the cell abundance (details in Methods). (b) Bar plot showing the relative differences of NPN uptake for the O-antigen expressing strain and LPS core mutants at low (0 to 1.563 μg/mL) and high (100 μg/mL) colistin concentrations. The zero line corresponds to the NPN uptake level of the parental strain. Error bars indicate standard errors of three replicates. (c) A table comparing expressed LPS components, electrokinetic properties, and outer membrane permeability of each strain. The ‘+’ / ‘–’ signs indicate the presence/absence of a corresponding LPS component. Percentages indicate the deviation of mean values relative to that of the parental strain.

## 4 Conclusion

Using microfluidic electrokinetic approaches, we have shown that *E. coli* carrying various LPS structures can be quickly distinguished by their linear electrokinetic mobility and cell polarizability. The presence of O-antigen in LPS leads to a decrease in the magnitude of linear electrokinetic mobility and an increased polarizability, which creates a less negatively charged but more polarizable cell surface. Correspondingly, O-antigen expression drives a strong resistance to colistin and low outer membrane permeability, which was observed in this study and elsewhere (Moffatt et al., 2019). In contrast, removing most of the core oligosaccharide by *waaC* deletion does not significantly affect cell electrokinetic mobility, but decreases cell polarizability and leaves cells highly susceptible to colistin. Truncating core oligosaccharides but retaining charged groups (e.g. phosphates) in the core by *waaF* deletion results in a strong negative electrokinetic mobility, a significantly increased polarizability, and an intermediate susceptibility and outer membrane permeability to colistin. A less drastic core alteration mediated by deletion of the heptosyltransferase *waaP* leads to a slightly more negative electrokinetic mobility, a strikingly higher polarizability, and a minor decrease in the bacterial resistance to colistin. Importantly, mutations in the *waa* region have been reported to cause reduced outer membrane protein abundance compared to wild-type cells, which may also drive changes in electrokinetic properties of bacteria. For instance, Δ*waaC* and Δ*waaF* mutations have been reported to cause the loss of flagella (Nakao et al., 2012) and a reduced abundance of outer membrane porins, OmpC and BamA (Halvorsen et al., 2021). In contrast, the Δ*waaP* mutant expresses higher levels of flagella and wild-type levels of OmpC and BamA (Nakao et al., 2012; Halvorsen et al., 2021). These changes to the cell surface may alter charge density and surface conductance of the soft layers around the cell and thus affect electrokinetic properties. Therefore, further proteomic analysis of membrane protein isolates will be needed to disentangle direct effects of LPS changes and porin abundance on these cell surface charge and dielectric properties.

The 3DiDEP and electrokinetic mobility measurement platforms could be scaled up to solve clinically important problems by leveraging recent technological advances. For example, mass production can be realized by microinjection molding with low cost and high reproducibility. Recent studies have shown that microinjection molding can create microstructures with a grooving resolution down to ∼4 *μ*m (Lu et al., 2019) and a roughness below 1 *μ*m (Liao et al., 2019), which aligns with the fabrication precision required for our 3DiDEP microchannel geometry (Supplementary figure 1). In addition, printed circuit board (PCB) can be used to add more complex electrical components (e.g. electrode pairs, sensors, and data acquisition ports) into microfluidic systems. For example, PCB has been bonded to PMMA microfluidics with an adhesive tape in previous work (Chang and You, 2019). PCB can also be integrated with highly parallel microfluidic circuits of 3DiDEP and electrokinetic mobility measurement units to further boost the throughput and scalability, where impedance measurement (Nakata et al., 2022) is an alternative method for label-free and rapid detection of 3DiDEP cell trapping.

In brief, our study finds relationships between *E. coli* LPS glycoforms, cell electrokinetic and dielectric properties, and outer membrane permeability. By depleting LPS biosynthetic enzymes WaaC, WaaF, and WaaP in *E. coli*, we show that core oligosaccharide changes drive changes in cell surface properties that can be detected in microfluidic electrokinetic devices. These results facilitate the development of high-throughput label-free platforms for screening or isolating bacteria based on their LPS structures using microfluidic electrokinetics. In this study, we used genetically induced LPS mutants as models to demonstrate the feasibility and efficacy of our approach, which paves the way for extended applications to identify and profile clinically relevant pathogens (e.g., *Salmonella enterica*) (Aldapa-Vega et al., 2019) that differ by a variety of LPS modifications, such as modifications in lipid A and O-antigen (Kim and Slauch, 1999; Raetz and Whitfield, 2002; Gulati et al., 2005; Meredith et al., 2007; Bogomolnaya et al., 2008) or capsule expression.

## 6 Acknowledgments

We acknowledge Nicole Chang at UCSB for providing *E. coli* strains, Noelani Kamelamela at MIT BioMicro Center for the equipment training, Dr. Jaehwan Kim for useful comments with potential applicability of the device. We are grateful to anonymous reviewers for their valuable comments.

## 7 Funding

The work was supported by the Department of Energy’s Genome Sciences Program grant SCW1039 and the National Science Foundation award number 1150615. HK was partly supported by the Kwanjeong Educational Foundation. SC and CRB were supported by the National Institutes of Health under Grants 1R01DE027850-01 and RM1 GM135102. TMH was supported by grant GM117930 from the National Institutes of Health.

## 8 Author contributions

QW and CRB conceived the study. QW, HK and SC designed and performed the experiments. TMH and CSH provided bacterial strains and guided the cell cultivation. QW, HK, and TMH interpreted the data. All authors wrote the paper. QW and HK have contributed equally to this work and share first authorship.

## 9 Conflict of interests

The authors declare that the research was conducted in the absence of any commercial or financial relationships that could be construed as a potential conflict of interest.

## Notes

### Competing Interest Statement

The authors have declared no competing interest.

### Summary of Updates

Author list and affiliations updated; New method section 2.10 added; Result sections 3.1 and 3.2 updated and new supplementary materials/figures added to clarify the effects of nonlinear electrophoresis and joule heating on cell motion.

