## Supplementary figures/tables/notes for "Leveraging microfluidic dielectrophoresis to distinguish compositional variations of lipopolysaccharide in *E. coli*"

### *Supplementary Material*

#### 1 Supplementary Figures and Tables

##### 1.1 Supplementary Figures

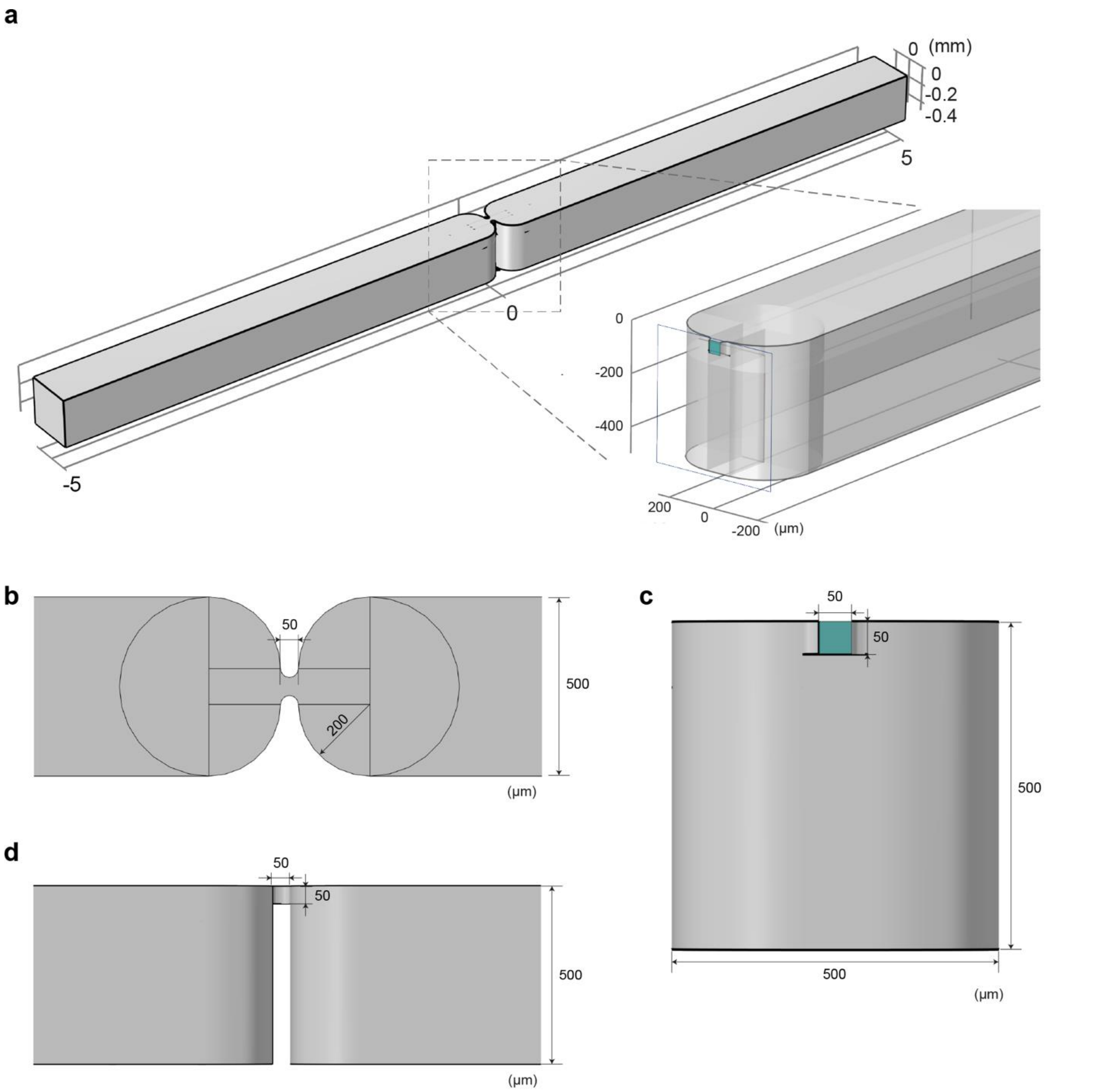


**Supplementary Figure 1.** Dimensions of the 3DiDEP microchannel. (a) An isometric view and a magnified cross-section view at the center of the 3DiDEP microchannel. (b) Top-, (c) front-, and (d) side-view of the microchannel. The cross-section plane at the center of the channel constricting region is highlighted in green. Electrodes were assembled by inserting platinum wires at both ends of the microchannel.

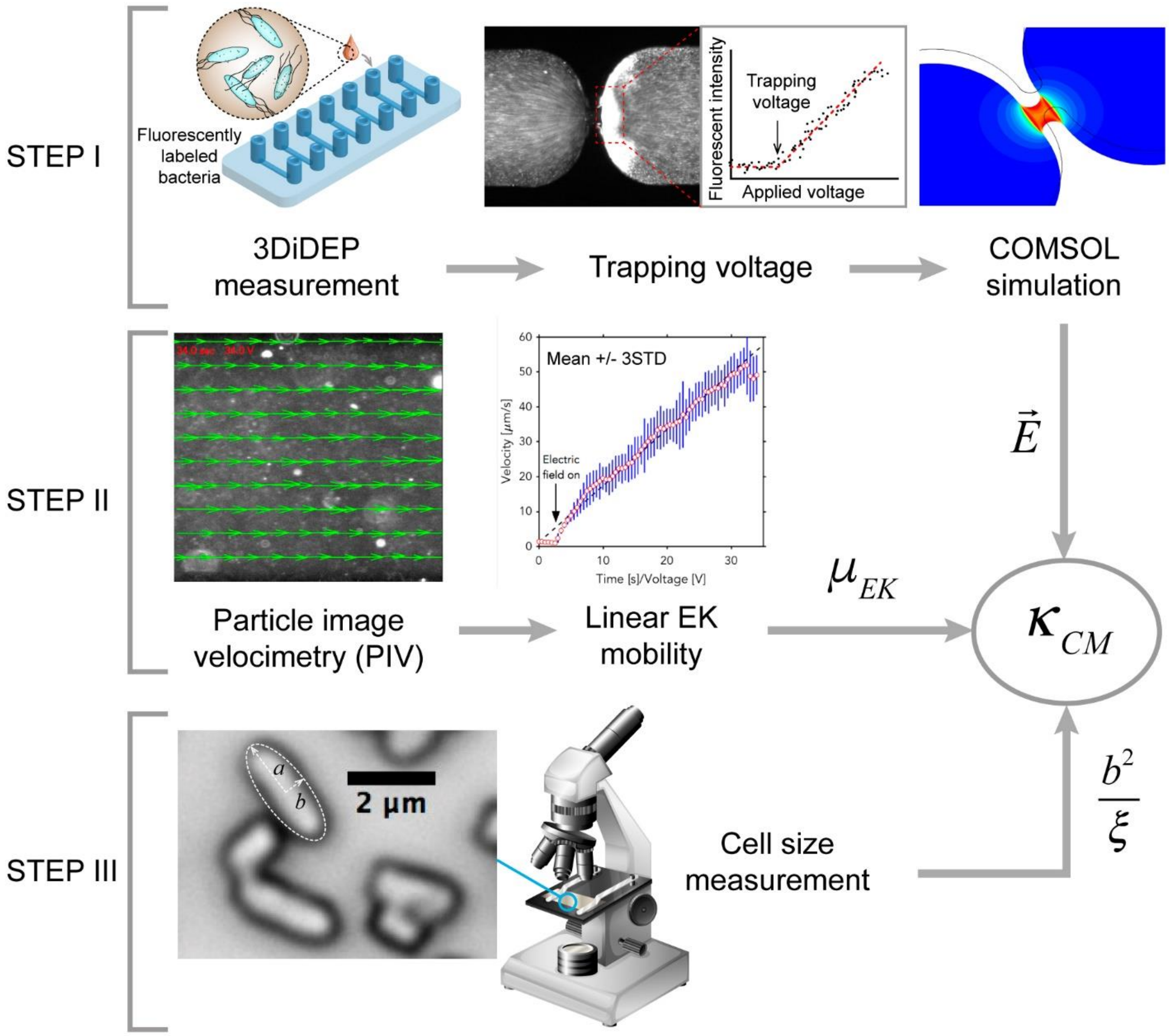


**Supplementary Figure 2.** Workflow for the derivation of cell polarizability (Clausius-Mossotti factor). STEP I, the measured trapping voltage required for the initiation of 3DiDEP cell immobilization was input as boundary conditions into a COMSOL 3D model to estimate the critical electric field. STEP II, particle image velocimetry (PIV) was used to monitor cell velocity changes in response to an increasing external electric field to determine the linear electrokinetic mobility. STEP III, an ellipsoidal fit was used to extract cell major (*a*) and minor (*b*) semi-axes and derive the corresponding Perrin friction coefficient from high-magnification images. Cell polarizability (Clausius-Mossotti factor) was estimated from the three parameters according to Equation S17 and S18. Adapted from (Wang, Q., 2018), copyright MIT Libraries.

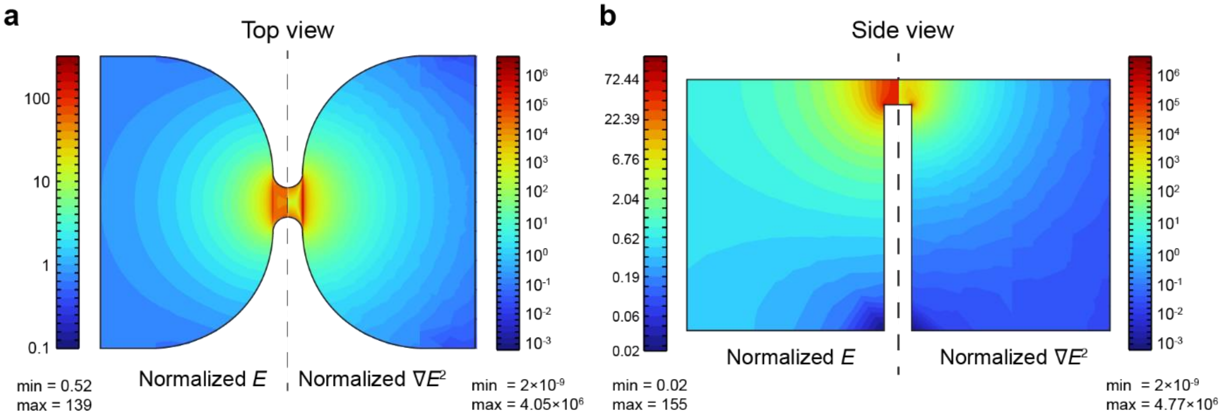


**Supplementary Figure 3.** Contour distributions of normalized electric fields ($E$, left) and the gradient of electric field squared ($\nabla E^2$, right). The electric field is normalized to the ratio of trapping voltage ($V_{trap}$) to the channel length ($L$). The ∇ operator is normalized to the inverse of the 3DiDEP constriction width ($1/W$). (a) A top view of the X-Y cross-section sitting at the floor of the 3DiDEP constriction (Z = -50 μm). (b) A side view of a X-Z cross-section at the center of the microchannel (Y = 0).

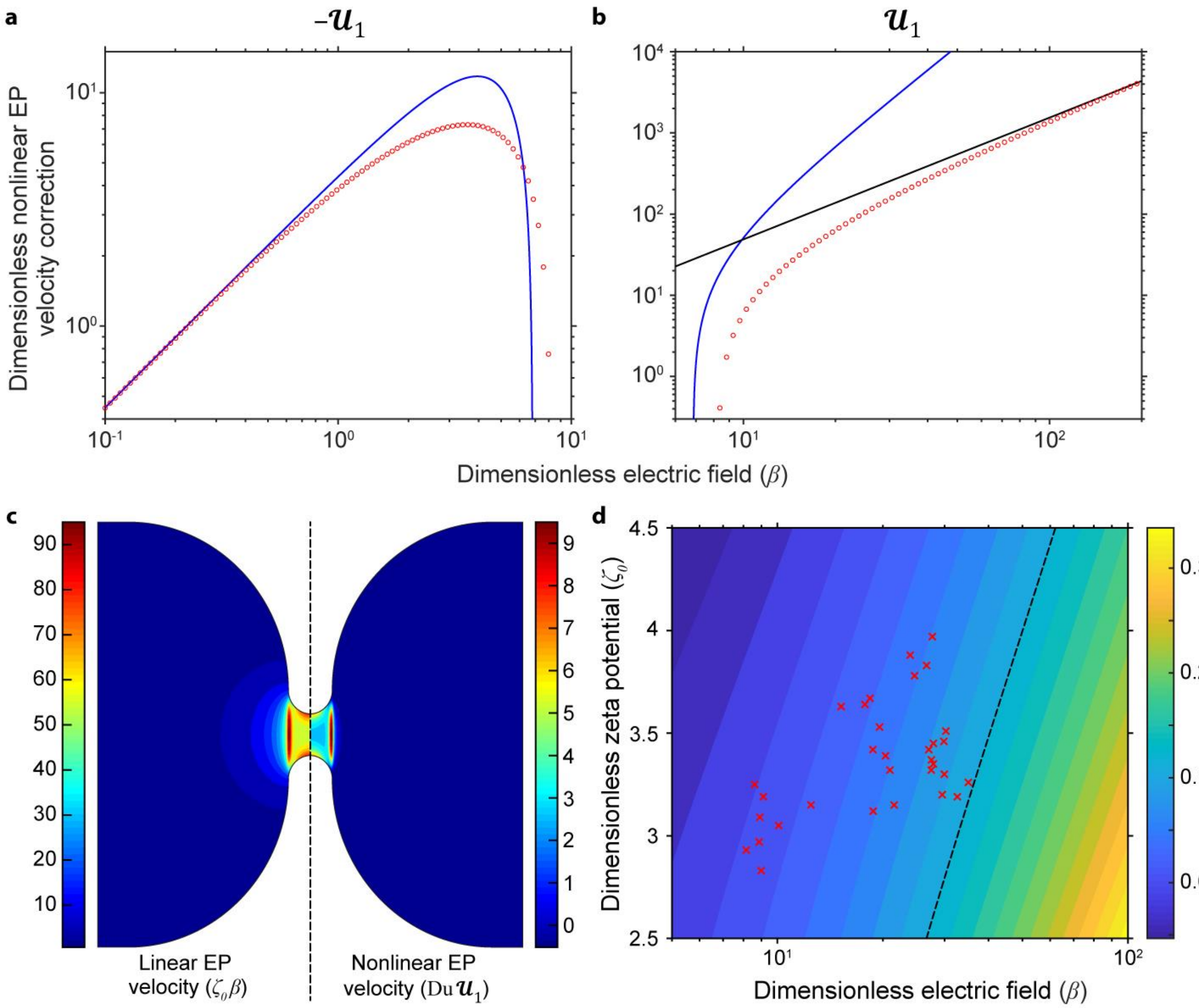


**Supplementary Figure 4.** The nonlinear electrophoretic (EP) velocity correction ($\mathcal{U}_1$) as a function of the dimensionless electric field ($\beta$) for a representative cell surface charge condition ($\zeta_0$ = 3.26, $\alpha$ = 0.282, $\dot{\alpha}$ = 0.059) measured for the $\Delta waaC$ mutant which corresponds to the highest trapping voltage. Due to the transition from negative to positive values, an estimation of $-\mathcal{U}_1$ (weak fields) and $\mathcal{U}_1$ (stronger fields) using the Schnitzer-Yariv model (red circles, Equation S4) are shown in (a) and (b), respectively. Also shown are the strong-field approximation (black line, Equation S11) and small-ion approximation (blue curves, Equation S12). (c) A comparison of linear (left) and nonlinear (right) dimensionless EP velocities at the highest measured trapping voltage (88.6 V, other conditions are consistent with panels a and b). (d) The ratio of nonlinear EP velocity to linear EP velocity as a function of the dimensionless zeta potential ($\zeta_0$) and dimensionless electric field ($\beta$). Our data (red crosses) show that the nonlinear EP velocity is less than 12% (dashed line) of their linear counterparts for all the tested conditions. The dimensionless parameters used for the analysis are explained in Supplementary Note 3.1 in detail.

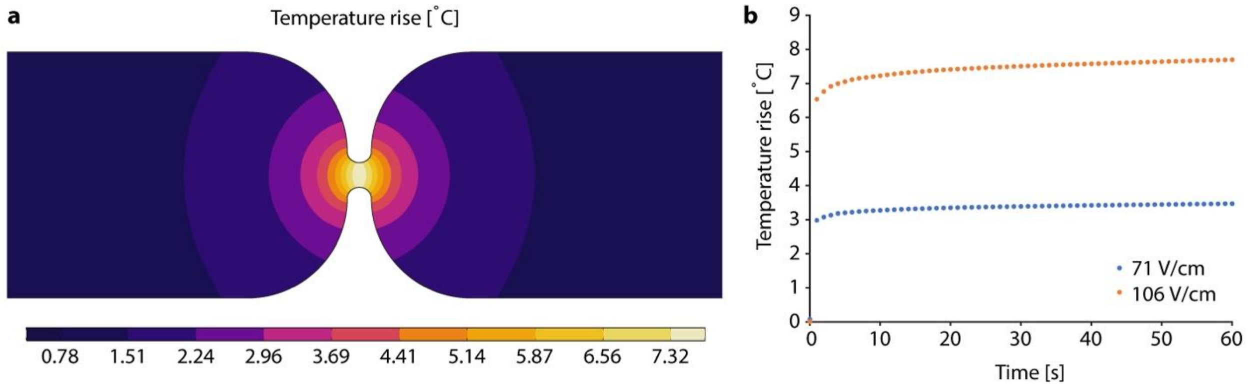


**Supplementary Figure 5.** Temperature rise near the 3DiDEP constriction 60 s after the application of a constant 106 V/cm electric field computed by a COMSOL simulation (Wang et al, 2017). (b) Maximum transient temperature rises within the microchannel in response to the application of a constant 71 V/cm (blue) and 106 V/cm (orange) electric field.

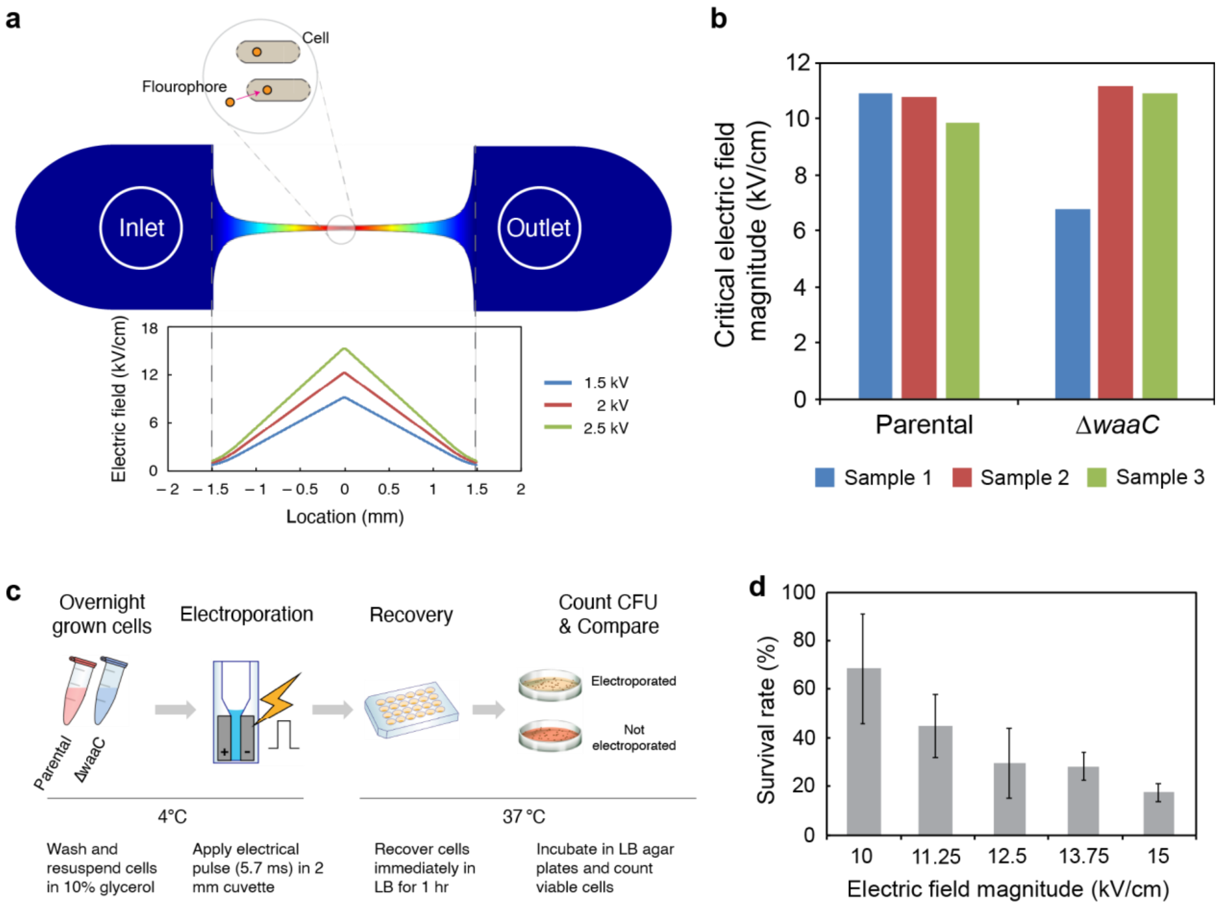


**Supplementary Figure 6.** (a) Schematic of a bilaterally converging microchannel used to measure the critical electric field ($E_{crit}$) that induces cell membrane permeabilization (Garcia et al., 2016, 2017), along with a 2D electric field distribution (red indicates strong values). Also shown are the linearly distributed electric fields along the channel centerline (bottom). Upon the application of an electric pulse, a section of the channel constriction region illuminates due to the SYTOX dye uptake by cells with enhanced membrane permeabilization, when the local electric field exceeds $E_{crit}$. (b) The critical electric field magnitudes ($E_{crit}$) measured for *E. coli* parental and Δ*waaC* mutant strains using the 2D microfluidic assay shown in (a). (c) Experimental workflow for measuring cell viability under strong electric fields using commercially available cuvettes. (d) Survival rates of the *E. coli* Δ*waaC* mutant strain measured by the ratio of CFU counts after and before the application of an electric field ranging from 10 to 15 kV cm$^{-1}$ using electroporation cuvettes. Details of the experimental procedure is described in Section 3.4.

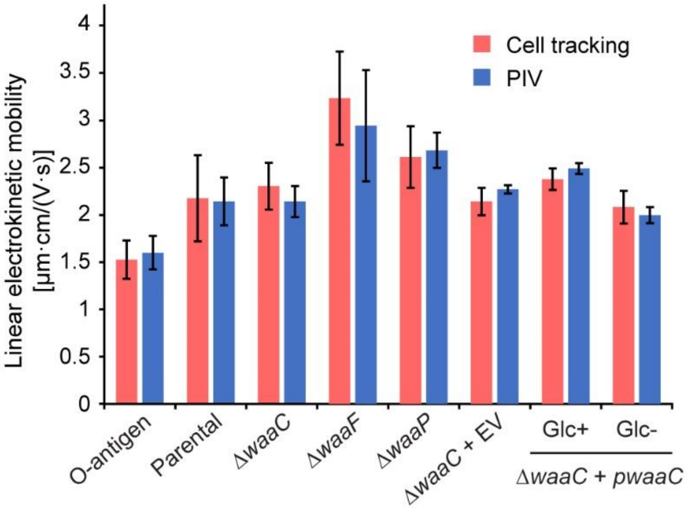


**Supplementary Figure 7.** Linear electrokinetic mobilities measured for each *E. coli* strain using cell tracking analysis (red, corresponding to main Figure 1b) or PIV analysis (blue) for cells in 3DiDEP experiments (corresponding to Figure 2a) show consistent results from the two approaches.

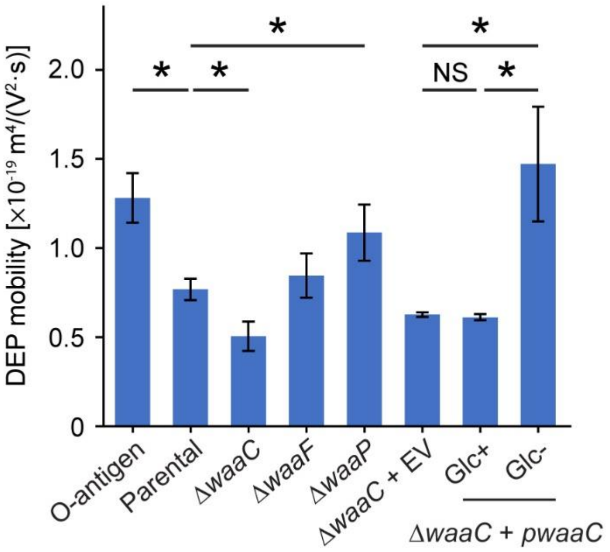


**Supplementary Figure 8.** DEP mobility measured for each strain. Asterisks indicate a statistical difference (*$p < 0.05$, NS: not significant).

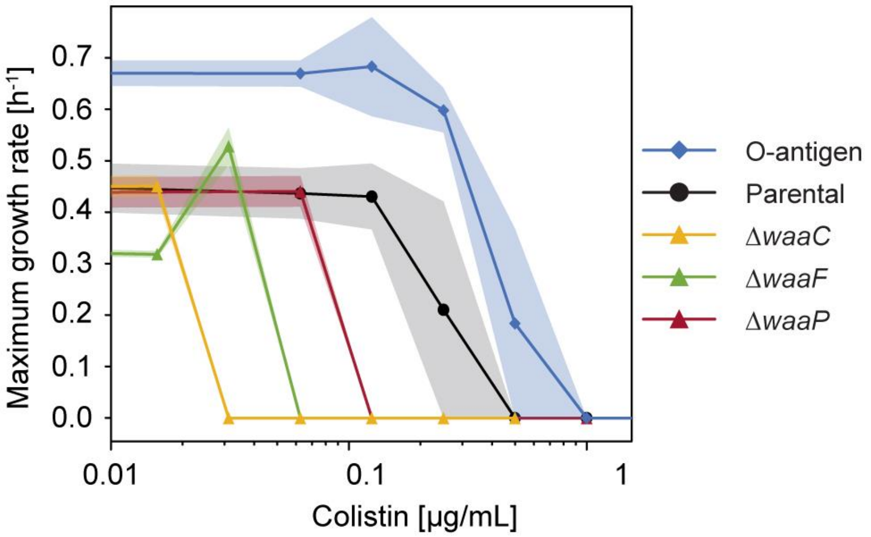


**Supplementary Figure 9.** Maximal growth rate of five *E. coli* strains from experiment testing for their colistin susceptibility. Growth rates were calculated from the abundance reads as described in Methods. All strains were grown in LB with 50 µg/mL streptomycin.

#### 1.2 Supplementary Table

**Supplementary Table 1.** Bacterial strains and plasmids

| **Strain** | **Genotype**[a] | **Description** | **Ref.** |
|---|---|---|---|
| DL5353 | *E. coli* K-12 W3110 pMF19, $Spm^R$ | W3110 expressing O16 polysaccharide (O-antigen). | Halvorsen et al. 2019; Feldman et al. 1999 |
| CH7175 | *E. coli* K-12 EPI100 Δ*wzb*, $Str^R$ | Parental stain used to make the *waa* mutants, expressing full LPS without O-antigen. | Halvorsen et al. 2019; Baba et al. 2006 |
| CH13815 | *E. coli* K-12 EPI100 Δ*wzb* Δ*waaC::kan*, $Str^R$ $Kan^R$ | EPI100 LPS lacking all sugars distal to $Kdo_2$-lipid A. | Halvorsen et al. 2019; |
| CH13816 | *E. coli* K-12 EPI100 Δ*wzb* Δ*waaF::kan*, $Str^R$ $Kan^R$ | EPI100 LPS truncated after HepI, lacking HepII and all moieties attached to it. | Halvorsen et al. 2019; |
| CH13817 | *E. coli* K-12 EPI100 Δ*wzb* Δ*waaP::kan*, $Str^R$ $Kan^R$ | EPI100 LPS lacking phosphates on HepI and HepII, and HepIII residues. | Halvorsen et al. 2019; |
| CH14472 | *E. coli* K-12 EPI100 Δ*wzb* Δ*waaC::kan* pCH450 empty vector, $Str^R$ $Kan^R$ $Tet^R$ | EPI100 Δ*waaC* strain transformed with an empty pCH450 plasmid. | This study |
| CH14473 | *E. coli* K-12 EPI100 Δ*wzb* Δ*waaC::kan* pCH450-*waaC*, $Str^R$ $Kan^R$ $Tet^R$ | Δ*waaC* complementary strain, whose *waaC* expression is suppressed in the presence of glucose. | This study |
| **Plasmid** | | | |
| pMF19 | Expresses the *wbbL* rhamnosyltransferase gene for O16 antigen synthesis, $Spm^R$ | | Halvorsen et al. 2019; Feldman et al. 1999 |
| pCH450 | pACYC184 derivative that carries *araC* and $P_{ara}$ promoter, $Tet^R$ | | Halvorsen et al. 2019; |

[a]Abbreviations: $Kan^R$, kanamycin resistant; $Spm^R$, spectinomycin resistant; $Str^R$, streptomycin resistant; $Tet^R$, tetracycline resistant.

**Supplementary Table 2.** Physical parameters used for nonlinear electrophoresis analysis

| Parameter | Symbol | Value | Unit |
|---|---|---|---|
| Medium viscosity | $\eta$ | 0.001 | Pa·s |
| Medium permittivity | $\varepsilon_m$ | 7.1E-10 | F/m |
| Thermal voltage | $\varphi_T$ | 25.69 | mV |
| Characteristic cell length | $a$ | 1 | μm |
| Diffusivity of $Na^+$ ions | $D_+$ | 1.33E-9 | $m^2/s$ |
| Diffusivity of $Cl^-$ ions | $D_-$ | 2.03E-9 | $m^2/s$ |
| Drag coefficient of $Na^+$ ions | $\alpha^+$ | 0.3407 | 1 |
| Drag coefficient of $Cl^-$ ions | $\alpha^-$ | 0.2232 | 1 |
| Debye length | $1/\kappa$ | 9.64E-9 | m |
| PMMA EOF mobility [a] | $\mu_{EOF}$ | 3.74E-8 | $m^2/(Vs)$ |

[a]EOF mobility of PMMA microchannels was measured following the protocol by Nohmi, M. & Santiago, J. (2008)

**Supplementary Table 3.** Estimated electrokinetic properties of studied bacteria strains

| Cell lines | Zeta potential (mV) [a] | Surface charge density ($mC/m^2$) [b] | Dimensionless zeta potential, $\zeta_0$ | Dimensionless electric field at cell trapping[c], $\beta$ | Du | Dimensionless nonlinear EP velocity, $\mathcal{U}_1$ | Ratio of nonlinear to linear EP, $R$ |
|---|---|---|---|---|---|---|---|
| O-antigen expressing | 75.1 ± 2.5 | 7.8 ± 0.4 | 2.974 ± 0.099 | 9.0 ± 0.7 | 0.0586 ± 0.0032 | 3.66 ± 2.22 | 0.008 ± 0.004 |
| Parental strain | 82.8 ± 3.6 | 9.2 ± 0.7 | 3.277 ± 0.141 | 20.0 ± 1.1 | 0.0692 ± 0.0052 | 57.93 ± 4.70 | 0.061 ± 0.003 |
| Δ*waaC* | 82.8 ± 2.3 | 9.2 ± 0.5 | 3.276 ± 0.091 | 30.8 ± 3.1 | 0.0691 ± 0.0035 | 155.88 ± 35.94 | 0.105 ± 0.013 |
| Δ*waaF* | 94.0 ± 8.3 | 11.8 ± 1.8 | 3.723 ± 0.328 | 24.9 ± 2.4 | 0.0884 ± 0.0138 | 74.99 ± 9.41 | 0.071 ± 0.005 |
| Δ*waaP* | 90.4 ± 2.6 | 10.8 ± 0.6 | 3.578 ± 0.104 | 17.9 ± 1.7 | 0.0812 ± 0.0044 | 35.06 ± 11.03 | 0.043 ± 0.010 |
| Δ*waaC* + EV | 84.6 ± 0.6 | 9.6 ± 0.1 | 3.348 ± 0.024 | 27.6 ± 0.2 | 0.0718 ± 0.0009 | 118.05 ± 2.25 | 0.092 ± 0.001 |
| Δ*waaC* + p*waaC* (Glc+) | 87.7 ± 0.8 | 10.2 ± 0.2 | 3.471 ± 0.031 | 29.3 ± 1.3 | 0.0766 ± 0.0013 | 126.43 ± 10.41 | 0.095 ± 0.004 |
| Δ*waaC* + p*waaC* (Glc−) | 80.7 ± 1.2 | 8.8 ± 0.2 | 3.196 ± 0.047 | 10.1 ± 2.1 | 0.0661 ± 0.0017 | 5.46 ± 8.69 | 0.009 ± 0.014 |

[a]Zeta potentials were estimated from measured linear electrokinetic mobilities ($\mu_{EK}$) at low electric fields (Figure 1d) and PMMA EOF mobility ($\mu_{EOF}$) using the Smoluchowski formula, $\mu_{EP} = \varepsilon_m \zeta / \eta$, where the magnitudes of $\mu_{EP}$ equals to the combination of $\mu_{EK}$ and $\mu_{EOF}$.

[b]Cell surface charge densities were estimated by the Gouy-Chapman relation, $\sigma^* = 2\varepsilon_m \kappa \varphi_T \sinh(\zeta / 2\varphi_T)$. Derivation of the dimensionless groups followed the equations in Section 3.1.

[c]The dimensionless electric field was evaluated using the maximum electric field generated near the channel constriction edge with the corresponding trapping voltage for each strain. Therefore, the estimated nonlinear EP velocities are the maximum values in the 3DiDEP microchannel at a given condition.

#### 2 Captions for Supplementary Videos

**Supplementary Video 1.** Representative videos showing 3DiDEP immobilization of the O-antigen expressing (top-left), parental (top-right), Δ*waaC* mutant (bottom-left), and Δ*waaF* mutant (bottom-right) *E. coli* strain. Cells were immobilized and gradually accumulated due to DEP near the microchannel constriction in response to the application of a 'linear sweep' DC potential difference (ramping rate, 1 V/s) across the microchannel (high potential applied to the right side). The red contours indicate the microchannel geometry, and the applied voltage is shown on the top-right corner of each panel. Blue arrows indicate the initiation of cell trapping. The threshold applied voltage at the onset of 3DiDEP immobilization is taken as the trapping voltage (Figure 2b) for the bacterial strain used. Scale bar, 100 μm. Video play speed is 5-folded increased (5 fps).

#### 3 Supplementary Notes

##### 3.1 Nonlinear electrokinetic effects in 3DiDEP

The Helmholtz-Smoluchowski equation for electrophoresis provides a linear relation between the electrophoretic (EP) velocity $\overrightarrow{u_{EP}}$ and applied electric field $\vec{E}$, $\overrightarrow{u_{EP}} = \mu_{EP}\vec{E}$, where the EP mobility, $\mu_{EP} = \varepsilon_m \zeta/\eta$, only depends on the dielectric permittivity $\varepsilon_m$ and viscosity $\eta$ of a symmetric electrolyte (with ionic valencies ± $Z$) and zeta potential $\zeta$ of the charged particle. However, Smoluchowski's theory assumes a weak electric field scenario, where the potential drop across the characteristic length of the particle $a$ is much smaller than the thermal voltage, $\varphi_T = k_B T/Ze$ (where $k_B T$ being the Boltzmann temperature and $e$ the elementary charge). When the dimensionless electric field $\beta = Ea/\varphi_T$ is much smaller than 1, the electrical double layer (EDL) can be considered as spherically symmetric and surrounded by an electroneutral bulk electrolyte. This EDL deforms as the applied electric field increases. When $\beta \gg 1$, the strong electric field breaks the symmetry of EDL and thereby creates tangential ionic currents (i.e., surface conduction). This surface conduction is intensified for highly charged particles and becomes significant for logarithmically large zeta potentials when $\zeta/\varphi_T \sim O(\ln(1/\delta))$, where $\delta$ is the dimensionless Debye length (Schnitzer, O. & Yariv, E., 2012). The non-uniform surface conduction over the curved particle surface necessitates compensating normal ionic fluxes across the EDL, which modifies the potential and ionic concentration distributions in the bulk electrolyte, giving rise to the nonlinear EP mobility that is no longer independent of the electric field strength.

Several prior studies of 2D insulated electrokinetic systems showed that nonlinear EP becomes increasingly significant as the applied electric field increases and counteracts the particle motion due to electroosmosis (EOF). As a result, the particle velocity due to the combination of EOF, linear, and nonlinear EP deviates from the linear Smoluchowski's prediction and drops to zero where particle trapping takes place (Cardenas-Benitez, B. et al., 2020, Antunez-Vela, S. et al., 2020, and Tottori, S. et al., 2019). Although these studies demonstrate good agreement between particle velocity measurements and a nonlinear electrophoresis model by Schnitzer and Yariv (Schnitzer, O. et al., 2013), cell trapping in our 3DiDEP system presents a distinctive physical scenario for the following three reasons.

1. The DEP force generated in these two-dimensional insulating constrictions is small due to the relatively small electric field gradient, which requires the use of large correction factors in numerical simulations to resemble the particle trapping observations in prior 2DiDEP studies (Hill, N. & Lapizco-Encinas, B. H, 2019 and Cardenas-Benitez, B. et al, 2020). In contrast, the DEP force at a 3DiDEP constriction is over 100 times stronger than the force estimated in

a corresponding 2D insulating constriction (with a geometry projected from 3DiDEP), because the DEP force is proportional to $\chi^2$, where $\chi$ is the constriction ratio (Braff, W. A. 2012). This is evident by the fact that the DEP force is strong enough to balance EP and EOF even at a low electric field range where nonlinear EP is negligible compared to linear EP (e.g., the O-antigen expressing strain in Supplementary Table 3). Therefore, the use of large correction factors is not necessary.

2. Many prior studies used a 2D microchannel made of polydimethylsiloxane (PDMS, with a strong negative zeta potential) to examine particles and bacterial cells with a low surface charge density (below 5 mC/m$^2$). In this case, EOF initially dominates over EP in the linear regime. As the applied electric field increases, nonlinear EP velocity increases beyond the linear approximation. Therefore, the combined linear and nonlinear EP counterbalancing EOF is the dominating mechanism of particle trapping (Cardenas-Benitez, B. et al, 2020). In contrast, we used 3DiDEP devices made of PMMA with surface priming using KOH. This leads to a lower EOF mobility (3.74 ± 0.34 μm·cm/Vs) in diluted PBS solutions (with a conductivity of 100 μS/cm, pH = 7). In this experimental condition, EP exceeds EOF and cell trapping is driven by the balance between positive DEP and the combination of EOF and EP.
3. Many prior studies reported nonlinear EP induced by moderate electric fields with $O(\beta) = 1 \sim 10$ (Cardenas-Benitez, B. et al, 2020 and Tottori, S. et al., 2019). In this regime, a weakly-nonlinear asymptotic theory proposed by Schnitzer and Yariv (Schnitzer, O. et al, 2013) has been used to model EP velocity. This model scales nonlinear EP velocity with $O(E^3)$ but is restricted to the limit of $\beta \leq O(1)$. In the present study, we observed a trapping voltage ranging from 22.6 to 88.6 V. This yields a $\beta$ as low as 0.09 ~ 0.36 in the large straight channel regions and it abruptly increases to a range of 9 ~ 35 at the 3DiDEP constriction, where the weakly-nonlinear asymptotic theory is no longer accurate. Therefore, we only need to estimate the contribution of nonlinear EP to particle motion for the strong electric field regime, $\beta \sim O(10)$.

Here we show that the contribution of nonlinear EP velocity is small compared to linear EP velocity in 3DiDEP systems when the applied voltage is lower than 90 V, which is the usual case for 3DiDEP cell trapping. To model nonlinear EP velocity, we use the semi-analytic solution for nonlinear EP velocity at arbitrary field strengths and small Dukhin number (Schnitzer, O. & Yariv, E. 2014). In what follows, a dimensionless formulation is given: the length variable is normalized by cell characteristic length, $a$ = 1 μm; the electric field $E$ by $\varphi_T/a$; the fluid velocity by $\varepsilon_m\varphi_T^2/\eta a$; and the stress variables by $\varepsilon_m\varphi_T^2/a^2$. The SY model provides the following expansion of dimensionless EP velocity $\mathcal{U}$:

$$\boldsymbol{\mathcal{U}} = \boldsymbol{\zeta_0\beta} + \mathrm{Du}\boldsymbol{\mathcal{U}_1} + \cdots \tag{S1}$$

where we target to derive an analytical expression of the correction term $\mathcal{U}_1$ in terms of given dimensionless parameters. In specific, $\zeta_0$ denotes a dimensionless zeta potential under the zero-Dukhin-number approximation, which gives

$$\boldsymbol{\sigma} = \mathbf{2}\sinh\left(\frac{\boldsymbol{\zeta_0}}{\mathbf{2}}\right) \tag{S2}$$

where $\sigma$ is the dimensionless group that normalizes cell surface charge density by $\varepsilon_m\kappa\varphi_T$, and $1/\kappa$ is the Debye length. We define the ionic drag coefficients $\alpha^+$ and $\alpha^-$ for cations and anions in the electrolyte, respectively, and the Dukhin number

$$\mathrm{Du} = (\mathbf{1} + \mathbf{2}\boldsymbol{\alpha}^{-})\boldsymbol{\delta\sigma} \tag{S3}$$

where $\delta = 1/(\kappa a)$ is the dimensionless Debye length. This implies that the first and second terms in Equation S1 represents linear and nonlinear contributions of EP velocity, respectively. For arbitrary electric fields where $\beta > O(1)$, $\mathcal{U}_1$ can be computed as

$$\boldsymbol{\mathcal{U}_1} = \left(\frac{1}{6\pi}\right)\left\{\boldsymbol{6\pi\beta}\frac{1-\grave{\alpha}\zeta_0-\alpha\zeta_0}{\alpha} + \boldsymbol{12\pi\beta}\left[\mathbf{4}\ln\cosh\left(\frac{\zeta_0}{4}\right) + \frac{1-\grave{\alpha}\zeta_0}{\alpha}\right]\mathrm{C}^{(1)}(\mathbf{1}) - \boldsymbol{18\pi\beta^2}\tanh\left(\frac{\zeta_0}{2}\right)\left[\mathrm{C}^{(0)}(\mathbf{1}) - \frac{1}{5}\mathrm{C}^{(2)}(\mathbf{1})\right] - \frac{36\pi(1-\grave{\alpha}\zeta_0)\beta^2}{5\alpha\zeta_0}\mathrm{C}^{(2)}(\mathbf{1}) + \frac{12\pi(1-\grave{\alpha}\zeta_0)\beta^2}{\alpha\zeta_0}\left\{\frac{9}{10}\mathrm{C}^{(2)}(\mathbf{1}) - \frac{3}{2}\mathrm{C}^{(0)}(\mathbf{1}) - \int_1^\infty\left[\boldsymbol{f}(\boldsymbol{r})\mathrm{C}^{(0)}(\boldsymbol{r}) + \boldsymbol{g}(\boldsymbol{r})\mathrm{C}^{(2)}(\boldsymbol{r})\right]\boldsymbol{dr}\right\}\right\} \tag{S4}$$

where

$$\boldsymbol{\alpha} = \frac{\alpha^+ + \alpha^-}{2}, \grave{\boldsymbol{\alpha}} = \frac{\alpha^+ - \alpha^-}{2} \tag{S5}$$

$$\boldsymbol{f}(\boldsymbol{r}) = \frac{d}{dr}\left(\boldsymbol{r}^2\frac{dA}{dr}\right), \boldsymbol{g}(\boldsymbol{r}) = \frac{1}{5}\left[\frac{d}{dr}\left(\boldsymbol{r}^2\frac{dB}{dr} - \boldsymbol{6B}\right)\right] \tag{S6}$$

in which the functions *A* and *B* are defined as

$$\boldsymbol{A}(\boldsymbol{r}) = \frac{1-r^2+4r^5}{4r^6}, \boldsymbol{B}(\boldsymbol{r}) = -\frac{1-5r^2-2r^3+2r^5}{4r^6} \tag{S7}$$

In Equation S4, $\mathrm{C}^{(0)}(\boldsymbol{r}), \mathrm{C}^{(1)}(\boldsymbol{r})$, and $\mathrm{C}^{(2)}(\boldsymbol{r})$ are the coefficients of the first three Legendre polynomials used to expend the ionic concentration *c* (scaled by the electrolyte ionic concentration)

$$\boldsymbol{c} = \boldsymbol{3\beta}\sum_{m=0}^{\infty}\mathrm{C}^{(m)}(\boldsymbol{r};\mathrm{Pe})\mathrm{P}^{(m)}(\cos\boldsymbol{\theta}) \tag{S8}$$

This leads to the computation of a set of coupled linear ordinary differential equations

$$\frac{d^2\mathrm{C}^{(n)}}{dr^2} + \frac{2}{r}\frac{d\mathrm{C}^{(n)}}{dr} - \frac{n(n+1)}{r^2}\mathrm{C}^{(n)}(\boldsymbol{r}) = \mathrm{Pe}\left\{\left(\frac{1}{r^3} - \mathbf{1}\right)\left[\frac{n}{2n-1}\frac{d\mathrm{C}^{(n-1)}}{dr} + \frac{n+1}{2n+3}\frac{d\mathrm{C}^{(n+1)}}{dr}\right] + \left(\frac{1}{r} + \frac{1}{2r^4}\right)\left[\frac{n(n-1)}{2n-1}\mathrm{C}^{(n-1)}(\boldsymbol{r}) - \frac{(n+1)(n+2)}{2n+3}\mathrm{C}^{(n+1)}(\boldsymbol{r})\right]\right\}, \boldsymbol{n} = \mathbf{0}, \mathbf{1}, \mathbf{2}\ \ldots \tag{S9}$$

where the effective Péclet number $\mathrm{Pe} = \alpha\zeta_0\beta$ represents the ratio of the cell surface tangential ionic advection to the normal ionic diffusion at $O(\mathrm{Du})$. The corresponding boundary conditions for Equation S9 is

$$\begin{cases} \mathrm{C}^{(n)} = \mathbf{0} \text{ at } \boldsymbol{r} \to \infty \\ \frac{d\mathrm{C}^{(n)}}{dr} = \boldsymbol{\delta_{n,1}} \text{ at } \boldsymbol{r} = \mathbf{1} \end{cases} \tag{S10}$$

where $\delta_{n,1}$ is the Kronecker delta function. For a large applied electric field with $\beta > O(10^3)$, Equation S4 can be simplified to a strong-field approximation:

$$\boldsymbol{\mathcal{U}_1} = \frac{2}{7}\sqrt{\frac{6}{\pi}}\left(\mathbf{9} - \mathbf{4}\sqrt{\mathbf{3}}\right)\left(\mathbf{1} - \dot{\boldsymbol{\alpha}}\boldsymbol{\zeta_0} + \boldsymbol{\alpha}\boldsymbol{\zeta_0}\tanh\left(\frac{\zeta_0}{2}\right)\right)\left(\frac{\beta}{\alpha\zeta_0}\right)^{3/2} \tag{S11}$$

This indicates that at strong fields, nonlinear EP velocity scales with $O(E^{3/2})$. The model also provides a small-ion approximation for the case of small Péclet numbers realized under moderate fields ($\beta \sim O(1)$) and small values of $\alpha$:

$$\boldsymbol{\mathcal{U}_1} = -\left[\mathbf{4}\ln\cosh\left(\frac{\zeta_0}{4}\right) + \boldsymbol{\zeta_0}\right]\boldsymbol{\beta} + \frac{2}{21}\boldsymbol{\beta}^{\mathbf{3}} \tag{S12}$$

We truncated the infinite series of coupled ordinary differential equations (Equation S9) to the term $n$ = 3 and calculated $C^{(0)}(r)$, $C^{(1)}(r)$ and $C^{(2)}(r)$ with $\beta$ ranging from 5 to 1000 and $\zeta_0$ ranging from 2 to 5 using MATLAB's bvp4c routine. Following Equations S4-S7, the nonlinear EP velocity correction, $\mathcal{U}_1$, was solved as a function of $\beta$ and $\zeta_0$. As a comparison, $\mathcal{U}_1$was also estimated using the strong-field approximation (Equation S11) and small-ion approximation (Equation S12).

In Supplementary Figure 4a and 4b, we take the condition of the $\Delta waaC$ mutant as an example to show the distribution of $\mathcal{U}_1$ as a function of $\beta$. The $\Delta waaC$ mutant requires the highest applied electric field among all the studied strains to initiate 3DiDEP trapping (Figure 2b). Therefore, trapping of this strain is expected to display the most significant nonlinear electrophoresis. We found that $\mathcal{U}_1$ is negative at weak electric fields, which is consistent with the first term in Equation S12, indicating that the first effect of surface conduction is to retard particle motion (Supplementary Figure 4a). When the dimensionless electric field $\beta$ exceeds 7, $\mathcal{U}_1$ transitions to positive values, with the particle velocity enhancement gradually converging to ~$O(\beta^{3/2})$ (Supplementary Figure 4b). This is consistent with the strong-field approximation (Equation S12), and in this regime, the action of the applied electric field on the ionic concentration distributions in the bulk electrolyte becomes dominant. This trend of $\mathcal{U}_1$ distribution agrees well with Schnitzer and Yariv's results (Schnitzer, O. & Yariv, E. 2014). We also imported 3D distributions of $\beta$ calculated at measured trapping voltages via COMSOL simulation into MATLAB and compared the corresponding distributions of linear and nonlinear EP velocities for each strain ($\zeta_0\, \beta$ and Du $\mathcal{U}_1$, respectively; Supplementary Figure 4c). To estimate $\zeta_0$ and Du, we converted the linear electrokinetic mobilities measured at low electric fields (Figure 1d) to zeta potential $\zeta$ through Smoluchowski's formula and estimated surface charge density values for each studied strain using the Gouy-Chapman relation, $\sigma^* = 2\varepsilon_m \kappa \varphi_T \sinh(\zeta/2\varphi_T)$ (see Supplementary Table 3). We show that even for the highest measured trapping voltage (e.g., 88.6 V) the contribution of nonlinear EP velocity is an order of magnitude smaller than linear EP velocity and is only effective in regions close to the edge of the insulating constriction (Supplementary Figure 4c). Combining data of $\zeta_0$, $\beta$, Du, and physical parameters listed in the Supplementary Table 2, we evaluated the effect of nonlinear EP by the ratio $R = \mathrm{Du}\mathcal{U}_1/\zeta_0\beta$ for all the studied strains (Supplementary Table 3) and for a range of $\zeta_0$ and $\beta$ (Supplementary Figure 4d). Our data show that linear EP dominates over nonlinear EP ($R \leq 0.11$) for all the tested 3DiDEP trapping conditions. In addition, for cells with a surface charge density above 6 mC/m$^2$ and a 3DiDEP trapping voltage below 253 V ($\beta < 100$), the magnitude of induced nonlinear EP velocity does not exceed 18% of linear EP velocity. In summary, due to the small effect of nonlinear EP at given experimental conditions, we decide to maintain the assumption that 3DiDEP cell trapping occurs when DEP balances linear EP and EOF.

#### 3.2 Derivation of the Clausius-Mossotti factor

Cell surface polarizability was assessed using the Clausius-Mossotti factor ($\kappa_{CM}$) according to the previously reported approach (Wang et al., 2019). During 3DiDEP, cells are immobilized in the vicinity of the microchannel constriction when DEP balances drag forces induced by the combination of electroosmosis and electrophoresis. The *E. coli* strains are rod shaped and can be modeled as ellipsoidal particles with semi-axes *a* > *b* = *c* (Supplementary Figure 2). The resulting Stokes' drag is expressed as

$$\overrightarrow{\boldsymbol{F_D}} = \boldsymbol{6\pi\xi\eta a}\left(\overrightarrow{\boldsymbol{u_f}} - \overrightarrow{\boldsymbol{u_c}}\right) \tag{S13}$$

where $\overrightarrow{u_f}$ and $\overrightarrow{u_c}$ are velocities of the background flow and of the cell (zero for immobilized cells), respectively, $\eta$ is the viscosity of the surrounding medium, and the Perrin friction factor (Koenig et al., 1975) is

$$\boldsymbol{\xi} = \sqrt{\boldsymbol{1-p^2}}/\ln\left[\left(\boldsymbol{1}+\sqrt{\boldsymbol{1-p^2}}\right)/\boldsymbol{p}\right] \tag{S14}$$

where *p* = *b*/*a* is the cell aspect ratio. As mentioned in the above section, the effect of nonlinear electrophoresis is significantly smaller than linear electrophoresis at given experimental conditions. Thus, the velocity of background electrokinetic flows is a combination of linear electrophoresis and electroosmosis, which can be considered proportional to the applied electric field, $\vec{E}$, as

$$\overrightarrow{\boldsymbol{u_f}} = \boldsymbol{\mu_{EK}}\vec{\boldsymbol{E}} \tag{S15}$$

where $\mu_{EK}$ is the combined linear electrokinetic mobility. The DEP force exerted on a bacterium by a DC electric field can be expressed as

$$\overrightarrow{\boldsymbol{F_{DEP}}} = \boldsymbol{2\pi ab^2\varepsilon_m\kappa_{CM}\nabla\vec{E}^2} \tag{S16}$$

where $\varepsilon_m$ is the permittivity of the surrounding medium. Given Equations S13 and S16, the criterion for 3DiDEP cell immobilization is

$$\boldsymbol{\mu_{EK}}\vec{\boldsymbol{E}}\cdot\vec{\boldsymbol{E}} + \boldsymbol{\mu_{DEP}}\left(\boldsymbol{\nabla}\vec{\boldsymbol{E}}^{\boldsymbol{2}}\right)\cdot\vec{\boldsymbol{E}} = \boldsymbol{0} \tag{S17}$$

where the DEP mobility, $\mu_{DEP}$, is specified as

$$\boldsymbol{\mu_{DEP}} = \frac{\boldsymbol{b^2\varepsilon_m\kappa_{CM}}}{\boldsymbol{3\eta\xi}} \tag{S18}$$

On the basis of Equation S17 and S18, DEP mobility and the Clausius-Mossotti factor can be estimated from the experimentally determined linear electrokinetic mobility (Figure 1d) and 3DiDEP trapping voltage (Figure 2b).

##### 3.3 Impact of nucleic acid staining on electrokinetic mobility measurement of bacterial cells

Because nucleic acid dyes are positively charged (Thermo Fisher Scientific Inc., 2014), an excessive dosage may shield the negative surface charges carried by LPS and impact the electrokinetic characterization of the cells. However, this effect has been proven to be highly dependent on the dye concentration. A previous study found that a high concentration of SYTO9 can alter the fatty acid composition in the cell membrane (Deng et al., 2020). However, this change became almost negligible when SYTO9 concentration was below 2.5 uM.

In this study, we used SYTO dyes at a much lower concentration of 5 nM. In addition, we used a cell concentration ($OD_{600}$ 0.3–0.6, equivalent to ~5 $\times$ $10^8$ cells/ml) much higher than those presented by Deng et al. (~$10^6$ cells/ml). This indicates that each cell was exposed to a dye concentration ~$10^4$

times less than the threshold concentration that can induce a physicochemical change in the cell membrane. Moreover, cells were repeatedly washed and resuspended in fresh DEP buffer after staining to further remove excessive dyes in the background medium. Therefore, we conclude that cell surface electrokinetic properties were not significantly influenced by SYTO BC at such a low concentration.

##### 3.4 Impact of Joule heating and strong electric fields on cell viability and membrane integrity during 3DiDEP

We next seek to determine whether high electric fields and the induced thermal effects in the 3DiDEP channel constriction lead to significant impact on bacterial cell viability and membrane integrity. Joule heating effects in this 3DiDEP system has been discussed in our previous work (Wang et al., 2017) where both analytical estimations and a 3D time-dependent numerical simulation performed using COMSOL were provided with supporting experimental validation. As shown in Supplementary Figure 5a, when a constant 106 V is applied across the one-centimeter-long channel for a minute, the maximum temperature rise is less than 8 ℃ at the center of the channel constriction. In addition, 85% of the maximum temperature rise is reached within the first second due to the quasi-steady state heat conduction within the fluid domain (Supplementary Figure 5b). In this study, we applied a DC voltage that increased from 5 to 100 V with a rate of 1 volt per second. Given that the highest trapping voltage was 88.6 V, each of the experiment was finished within 85 seconds. A trapping voltage of 100 V corresponds to an RMS constant voltage of 59 V, indicating that the maximum temperature rise is below 3 ℃. This temperature change is much smaller than the temperature difference between the cell culture condition (37 ℃) and the ambient temperature (20 ℃). Therefore, we conclude that joule heating does not significantly affect cell viability during 3DiDEP.

In addition to thermal effects, the application of strong electric fields may affect cell membrane permeability and cell viability. Many studies have reported that the transmembrane potential induced by the application of external electric fields can create reversible nanoscale defects in the cell membrane that allows for intracellular transport of foreign molecules (e.g. plasmids or drugs). Stronger electric fields can even lead to irreversible cell death. Here we show that the maximum electric field applied during 3DiDEP is below the threshold electric field that causes reversible cell membrane permeabilization and much smaller than the threshold value that leads to cell death. We present a new data testing for the critical electric field magnitude, defined as the minimum value that induces cell membrane disruption. To measure cell responses to a varying voltage more precisely, we set up two assays using a previously developed two-dimensional microchannel and commercially available high-voltage electroporation cuvettes.

First, to determine the critical electric field for reversible cell membrane permeabilization, we used a previously reported microfluidic device (Garcia et al., 2016, 2017), which contains a bilaterally converging channel that creates a linearly distributed electric field upon the application of an electric pulse (Supplementary Figure 6a). In brief, cells were washed with 200X diluted PBS to obtain a more compromised membrane and loaded into the channel in the presence of a SYTOX® Green nucleic acid stain (Life Technologies, Grand Island, NY). When an electric pulse (MicroPulser™) was applied, the SYTOX dye fluoresces and illuminates the location in the channel (highlighting the corresponding electric fields) where cell membrane permeabilization was induced (Supplementary Figure 6a). Our statistical analysis comparing fluorescent intensity change before and after the electric pulse application show that *E. coli* parental and *waaC* deficient strains did not exhibit significant cell membrane permeabilization in response to an external electric field below 10.5 and

9.5 kV/cm (Supplementary Figure 6b), respectively. These measured critical electric fields are higher than the maximum electric field (2~8 kV/cm) exposed by the cells when a trapping voltage was applied for all the studied strains, indicating that the electric field induced membrane permeabilization is insignificant during 3DiDEP in this study.

In addition, we measured cell survival rates in response to a variety of electric field intensities using commercially available electroporation cuvettes (Supplementary Figure 6c), following a standard electroporation protocol provided elsewhere (Garcia et al., 2016, 2017; Huang et al., 2022). In brief, Δ*waaC* mutant cells were washed twice with 10% glycerol (centrifuged at 3500 rpm for 5 min followed by 8000 rpm for 10 min at 4 ℃). The resulting cell suspension was loaded in a 2 mm-width electroporation cuvette, and an exponentially decaying electric pulse was applied with an amplitude ranging from 2 to 3 kV using MicroPulser™ (Bio-Rad, Hercules, CA) (duration = 6 ms; decaying constant = 6 ms). Before and after pulsing, the cell abundance was measured by CFU counting on LB agar plates supplemented with appropriate antibiotics. A survival rate was calculated by the ratio between CFUs counted before and after the pulse. We chose to perform electroporation using the Δ*waaC* mutant strain because it has the most LPS core truncated and exhibited the most vulnerable membrane barrier function among the studies strains (Figure 4a and 5). Our results show that ~68% of the cells survived and successfully recovered from an electric field of 10 kV/cm (Supplementary Figure 6d). Given that the glycerol treatment makes the cell membrane more electrocompetent then that of the cells used in the current study, and that the 3DiDEP induced maximum electric fields are lower than 10 kV/cm, we conclude that 3DiDEP did not significantly affect cell viability in this study.

In summary, an upper-bound estimation of temperature rise during 3DiDEP is below 3 ℃, suggesting that the Joule heating effects on cell viability is small. In addition, the critical electric field to initiate cell membrane disruption is 1–2 folds higher than those estimated at our 3DiDEP microchannel constriction sites at a given trapping voltage, suggesting that the effects of induced transmembrane potential on cell viability is also small in this study.
